# Missing microbial eukaryotes and misleading meta-omic conclusions

**DOI:** 10.1101/2023.07.30.551153

**Authors:** Arianna I. Krinos, Margaret Mars Brisbin, Sarah K. Hu, Natalie R. Cohen, Tatiana A. Rynearson, Michael J. Follows, Frederik Schulz, Harriet Alexander

## Abstract

Meta-omics has become commonplace in the study of microbial eukaryotes. The explosion of available data has inspired large-scale analyses, including species or taxonomic group distribution mapping, gene catalog construction, and inference on the functional roles and activities of microbial eukaryotes *in situ*. However, genome and transcriptome databases are prone to misannotation biases, and meta-omic inventories may have no recoverable taxonomic annotation for more than half of assembled contigs or predicted proteins. Direct mapping solely to organisms of interest might introduce a problematic misattribution bias, while full databases can annotate any cataloged organism but may be imbalanced between taxa. Here, we explore the potential pitfalls of common approaches to taxonomic annotation of protistan meta-omic datasets. We argue that ongoing curation of genetic resources is critical in accurately annotating protists *in situ* in meta-omic datasets. Moreover, we propose that precise taxonomic annotation of meta-omic data is a clustering problem rather than a feasible alignment problem. We show that taxonomic membership of sequence clusters demonstrates more accurate estimated community composition than returning exact sequence labels, and overlap between clusters can address database shortcomings. Clustering approaches can be applied to diverse environments while continuing to exploit the wealth of annotation data collated in databases, and database selection and evaluation is a critical part of correctly annotating protistan taxonomy in environmental datasets. We re-analyze three environmental datasets at three levels of taxonomic hierarchy in order to illustrate the critical importance of both database completeness and curation in enabling accurate environmental interpretation.

## Main

Protists (microbial eukaryotes) are ubiquitous and essential organisms that provide multifarious ecosystem services, ranging from interactions with other microbes to impact on global biogeochemical cycles^1–5^. Protists have complex ecosystem roles and morphology, and often bridge seemingly disparate scales of interactions, which makes them difficult to visually differentiate yet critical to census for a complete understanding of ecosystem ecology^1, 3, 4^.

Molecular surveys of microbial communities have allowed researchers to characterize taxonomic diversity without microscopy or imaging. Computational approaches are used to assess the taxonomic composition of metagenomic or metatranscriptomic samples. These may include k-mer profiling of raw reads^6–8^, direct recruitment of raw reads from the meta-omic (community-level) sequencing sample to a reference or set of references of interest (e.g. genome, transcriptome, metagenome-assembled genome (MAG), or single-amplified genome (SAG))^9–12^, identification and recovery of well-known marker genes (e.g., 18S rRNA) from meta-omic raw reads or from assembled contigs, followed by phylogenetic alignment and within-sample quantification^13–16^, or sequence search of assembled contigs to a database, using match quality and percentage identity cutoffs to assign best-available level of confidence to taxonomic annotation of genes^17–21^. Computational approaches to assign taxonomic identities range in the scale over which they can be applied (Supplementary Figure 1).

All annotation methods share a reliance on databases containing labeled sequences from past studies (“reference sequences”), some of which may carry study-specific features. Environmental microeukaryote meta-omic studies often rely on annotations from transcriptomes of cultured representatives of protists^12, 25–28^, and are therefore representative of conditions or treatments specific to an experiment. Though transcriptomes represent a fraction of the genome, they are more readily available than genomes due to the high time and monetary cost of sequencing the large repetitive and intergenic regions common to eukaryotes^29^. Because genetic data are often collected when cell densities and expression levels are high, expression profiles may be most similar to when the microorganism is dominant and abundant in the field. Moreover, reference datasets that include different cell life cycle stages and environmental conditions would be ideal to link taxonomic identity to functional role but are not always available^30^.

Here, we highlight three vignettes that span three scales of taxonomic hierarchy (genus, family, and phylum) and explore how database-informed taxonomic annotation of assembled predicted proteins may be impacted by database composition. We then apply a two-stage clustering technique that includes unsupervised clustering to each of these data vignettes to demonstrate how such a method may maximize accurate classification of sequences at all three taxonomic scales even as databases grow. We propose that clustering is essential for understanding the limit of our ability to taxonomically annotate *de novo* assembled sequences. Our method re-poses taxonomic annotation as a clustering problem and can be used to improve characterization of community composition at multiple levels of taxonomy, even if a complete database is unavailable.

### **Genus**: Genetic differentiation between species complicates accurate identification of genus-level community composition

Species in the haptophyte genus *Phaeocystis* are genetically related, yet have distinctive geographic distributions and morphologies. *Phaeocystis antarctica*, *P. globosa*, and *P. pouchetii* are often cold-adapted and form colonies and large blooms at high latitudes (“colony-formers”), while *P. cordata* and *P. jahnii* are found at mid-latitudes and do not form colonies (“free-livers”)^31–34^. We re-analyzed *Tara* Oceans metagenomic samples from the Mediterranean Sea and the Southern Ocean, assembling contigs and then annotating using standard lowest common ancestor (LCA) algorithm against three modified MMETSP and MarRef databases containing: 1) all *Phaeocystis* references (both colony-formers and free-livers), 2) only the colony-formers, and 3) only the free-livers; all databases contained non-*Phaeocystis* taxa. Given that all three databases contain *Phaeocystis* representatives to the genus level, our expectation was that all three databases would differentiate *Phaeocystis* at the genus level. In the Southern Ocean where large blooms of *P. antarctica* are observed, 79.0% of the total *Phaeocystis* sequences identified with a combined database were identified using the colony-former database, whereas only 11.3% of the *Phaeocystis* sequences were identified using the free-liver database (Figure 2). In the Mediterranean Sea where free-livers dominate, 58.8% of *Phaeocystis* sequences were identified using the free-liver database as compared to 39.9% with the colony-former database (Figure 2). This implies that the presence of biogeographically distinct species ecotypes in our databases complicated reliable identification of expected taxa – ecotypes that have not been added to the database may be entirely missed.

**Figure 2.**
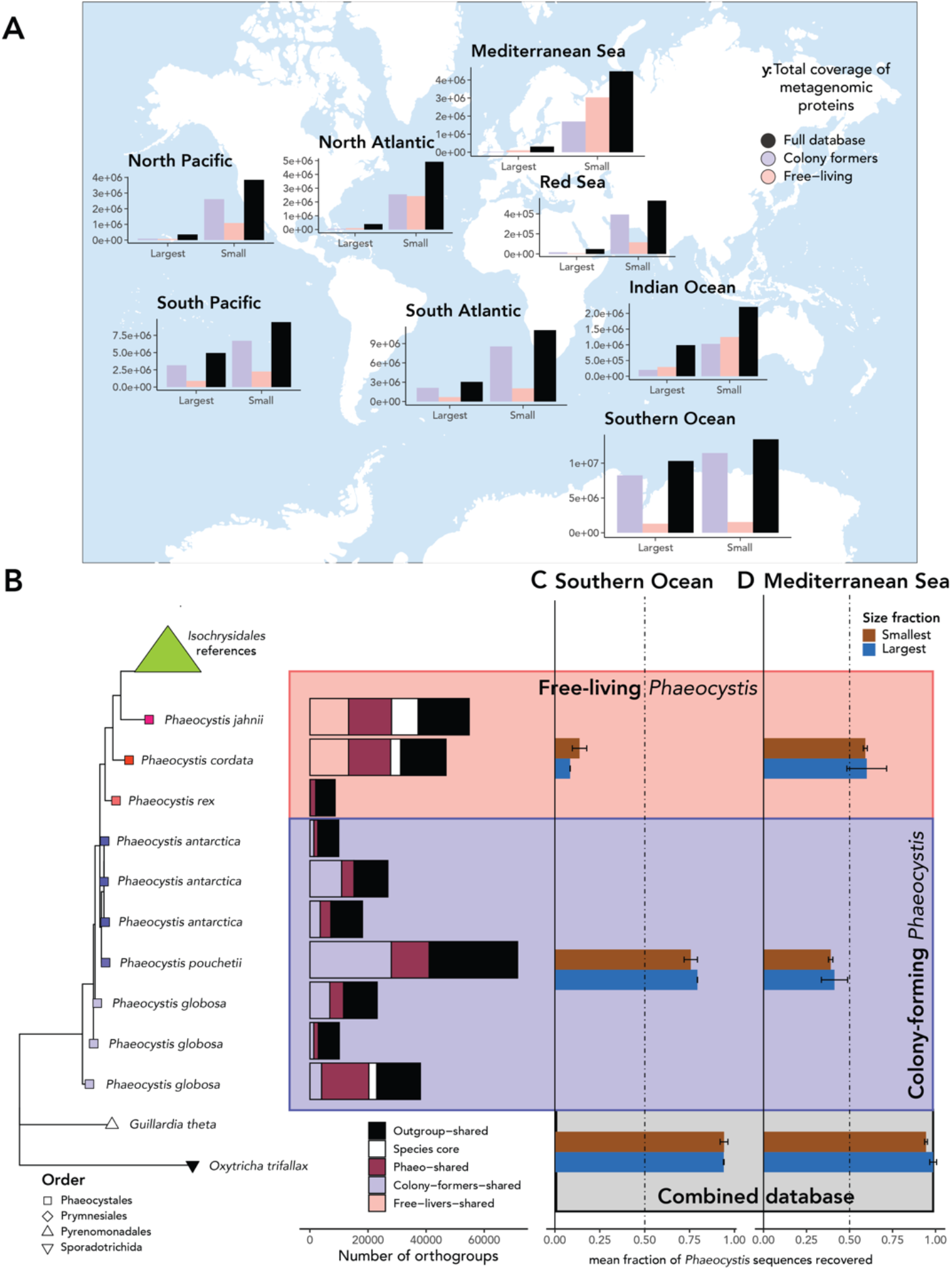
Effect of different species-level references on the success of genus-level identification of *Phaeocystis*. A: Abundance of metagenomic proteins in each ocean basin coassembled from the *Tara* Oceans dataset annotated to be *Phaeocystis* by a combined database of the colony-forming references (left in each group; purple), a combined database of the free-living references (middle in each group; pink), a combined database of all *Phaeocystis* references (right in each group; black). Each group of bars represents either the large or the small size fraction samples. B: Phylogenetic tree of *Phaeocystis* references and genomic and transcriptomic outgroups. The bars to the right of the tree show the total number of orthogroups in each species that are a, pink or lavender: shared by other members of the same ecotype (colony-former or free-liver), b, maroon: shared among multiple *Phaeocystis* species regardless of ecotype, or c, white: present only within one species. C: Percentage of sequences from the coassembly from the Southern Ocean *Tara* Oceans samples annotated to be *Phaeocystis* by any of the databases that were annotated as *Phaeocystis* using (top group of two bars) a combined reference database containing all of the free-living *Phaeocystis* references, (middle group of bars) a combined reference database containing all of the colony-forming *Phaeocystis* references, (bottom group of bars) a combined reference database containing all *Phaeocystis* references. The top bar in each group (brown) corresponds to the smallest *Tara* Oceans size fraction, while the bottom bar in each group (blue) corresponds to the largest *Tara* Oceans size fraction. D: Identical to Panel C, but for the *Tara* Oceans samples from the Mediterranean Sea.

### **Family**: Database imbalance limits phylogenetic resolution in closely-related diatom taxa

Taxonomic annotations can also yield unpredicted results within a taxonomic group. When a large number of reference sequences belong to one family, but none or only a few references belong to another, this imbalanced database representation may alter annotation recovery unexpectedly. We explored this phenomenon using metatranscriptomic data from a 2012 survey^26^ paired with associated microscopic cell counts (University of Rhode Island Long-Term Plankton Time Series; https://web.uri.edu/gso/research/plankton/data/). We focus our analysis on diatoms, a group that is well-represented in reference databases (266 transcriptomes in MMETSP), but has uneven representation across families (Anderson-Darling Test against uniform distribution: An=70.221; p=1.3e-5). The diatom *Dactyliosolen fragilissimus* (family *Rhizosoleniaceae*) constituted over 38-60% of the cells counted using light microscopy in 3 of 4 sampled weeks (Figure 3A). However, it was not consistently identified in the metatranscriptomes (<1% of species-level annotations)^26, 35^, despite the observed species being present in the reference database (Marine Microbial Eukaryote Transcriptome Sequencing Project (MMETSP)^27, 29, 36^. Four other *Rhizosoleniaceae* are also included in the MMETSP database^29^, yet the family constituted just 0.5-4.3% of family-level annotations and 0.1-0.7% of total sequence abundance. By contrast, the diatom family *Skeletonemataceae* represented as much as 95% of microscopy counts in one sample, and given the availability of isolates from Narragansett Bay in the database, it was well-annotated in the metatranscriptomes (Figure 3A). *Cerataulina pelagica* (family *Hemiaulaceae*) was also abundant in the microscopy data. Counterintuitively, while not present within the MMETSP database, contigs in the metatranscriptome were consistently annotated as belonging to *Hemiaulaceae* using a single related reference (*Eucampia antarctica*; Figure 3A). The outcomes of low database taxonomic resolution were incongruent between taxa: though both missing taxa of *Hemiaulaceae* and *Rhizosoleniaceae* had a member of the same family available in the database (Figure 3B), only *Hemiaulaceae* yielded annotations at the expected taxonomic resolution. Critically, this implies that taxonomic coverage alone often does not lead to accurate phylogenetic labels.

**Figure 3.**
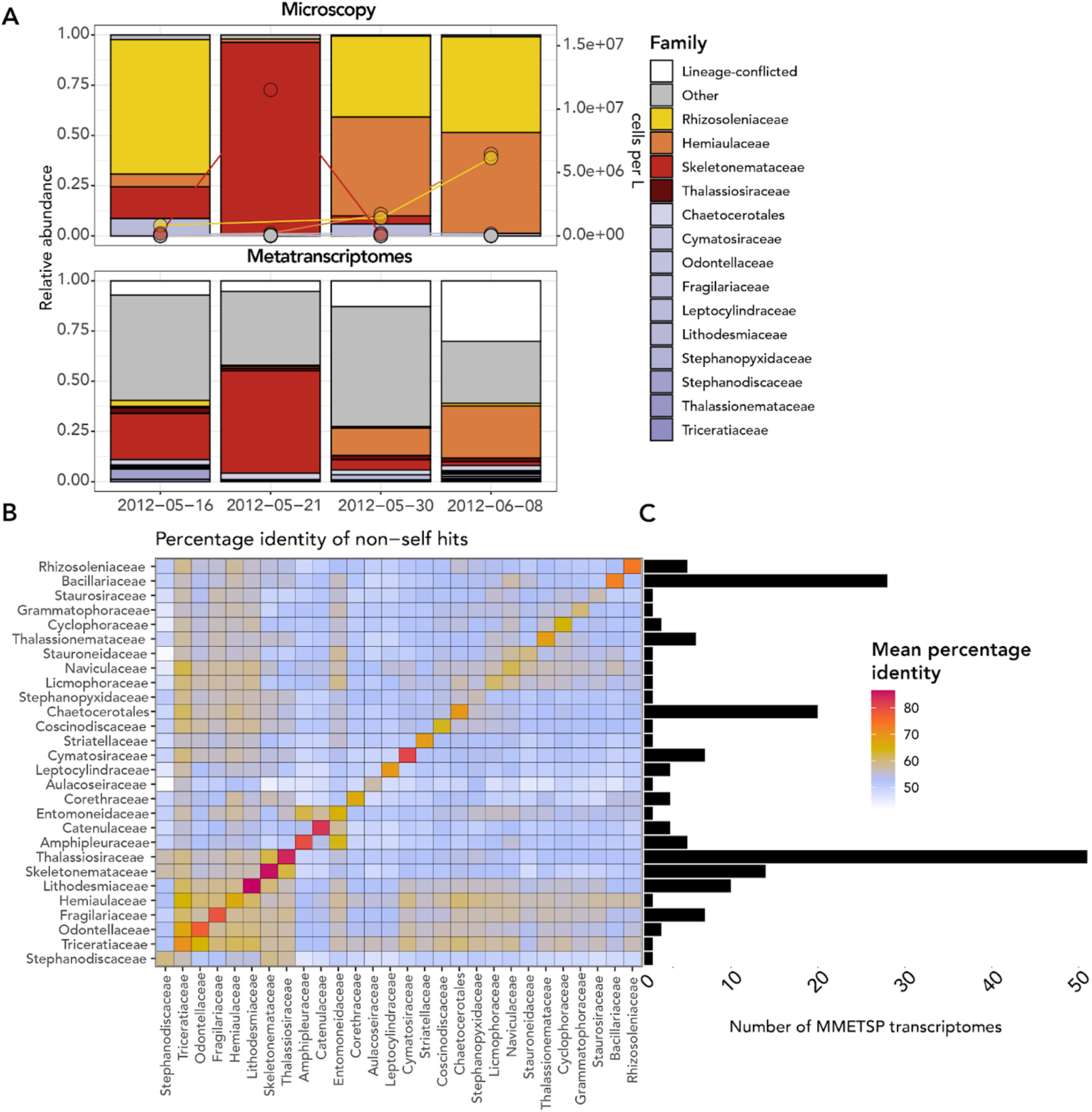
The effect of database composition on annotation of diatoms. A: Community composition of diatoms in Narragansett Bay based on light microscopy counts (top) compared to their metatranscriptomic activity (bottom). Lineage-conflicted refers to predicted proteins that were annotated as belonging to class Bacillariophyta, but had a conflict at the family level. “Other” refers to diatom families with associated TPM of less than 1,000. Circles (top) indicate cells per L (right y-axis). B: Mean percentage identity of non-self hits meeting a minimum bitscore value threshold (>=50) for diatom families represented in the MMETSP. The bars to the right of the plot indicate the total number of transcriptomes contained in the MMETSP for each family.

### **Phylum**: Broad-rank absence from databases leads to inaccurate community composition estimates

Sequence representation across major lineages in the eukaryotic tree of life is variable^1, 37^. We explored the impact of removing one eukaryotic lineage from a reference database on the predicted taxonomy of metatranscriptomes. Data from the North Atlantic along a transect from Woods Hole Oceanographic Institution (WHOI) to the Bermuda Atlantic Time Series (BATS) station (“BATS transect”) were annotated using a database lacking radiolarians (phylum Retaria). This left 42,736 putative radiolarian proteins unannotated and 46,283 annotated as different phyla across diverse lineages (Figure 4A-C). Adding radiolarians (see Online Methods) to the database impacted not only the total sequences labeled but also changed assigned annotations of existing taxa, highlighting how database incompleteness impairs community interpretation via both missing and incorrect annotations. Further, of 1,021,229 (8.6%) ORFs that were annotated at the domain–but not the phylum–level (“lineage-conflicted”), 95.8% were assigned a functional annotation, a higher rate than likelihood of functional annotation among all ORFs (45.8%). This suggests that highly conserved proteins will be left out of lineage-specific analysis because they tend to be taxonomically ambiguous (Figure 4D).

**Figure 4.**
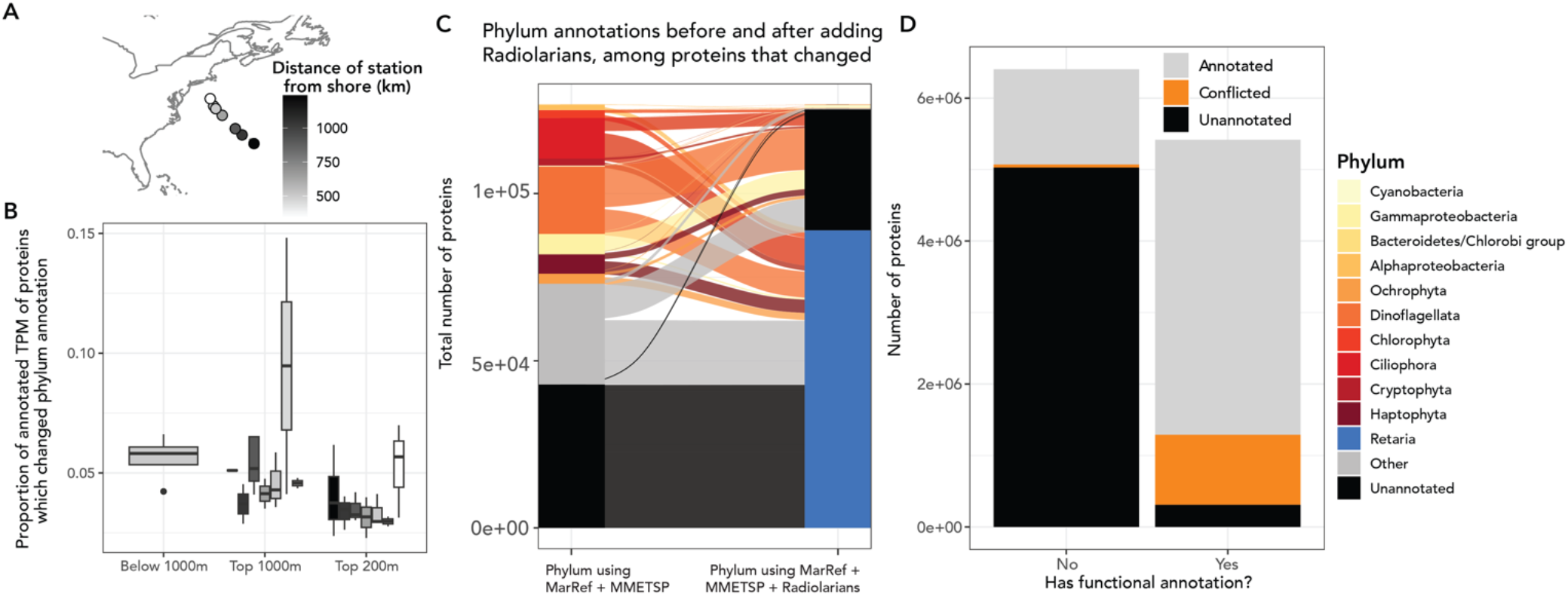
Effect of removing Radiolarian sequences from the database on the annotation of metatranscriptomic samples from the North Atlantic Ocean. A: Map of the BATS transect colored by the distance of each sample from the shore in kilometers. B: Fraction of annotated scaled abundance of proteins that changed annotation before and after the radiolarian sequences were added, grouped by depth. C: Among sequences that changed annotations, comparison of their annotation without radiolarian sequences (left axis) to with radiolarian sequences (right axis). In both cases the database contained the MMETSP and MarRef2 databases. While the majority category of putative Radiolarian sequences was those previously unannotated at the phylum level, some were previously classified as other phyla. Some phylum-level annotations were lost due to conflicts with added radiolarian sequences. D: Comparison of the number of proteins that were taxonomically annotated (“Annotated”), taxonomically unannotated (“Unannotated”), or had conflicting taxonomy (“Conflicted”) according to whether they were also functionally annotated.

**Figure 5.**
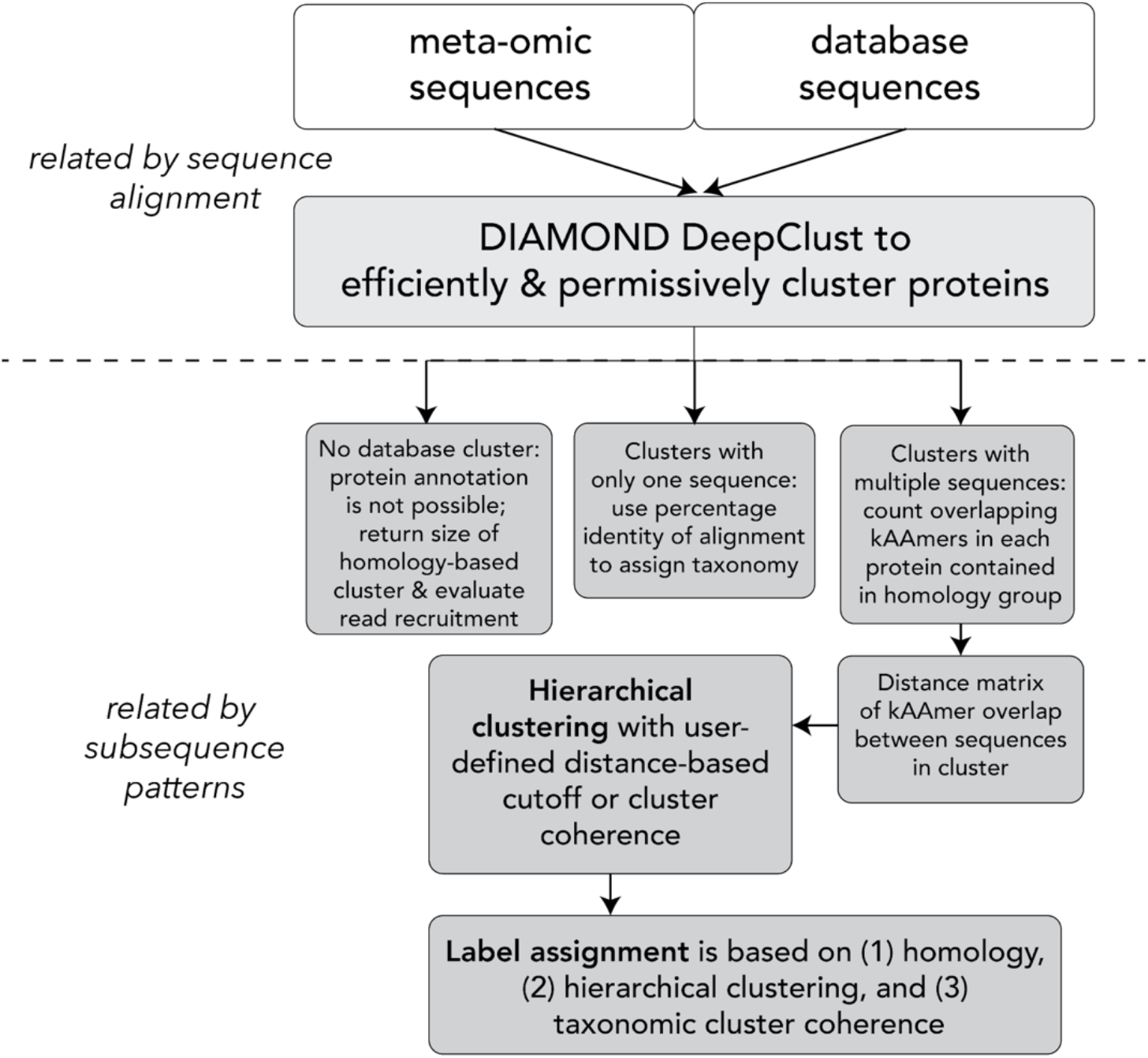
Schematic diagram of the tax-aliquots two-stage clustering workflow. The workflow is intended to be used alongside the LCA algorithm to detect ambiguity in taxonomic assignment and identify possible taxonomic annotations of sequences which cannot be annotated using the short alignment method. By assessing similarity using subsequence patterns over the entire sequence length, tax-aliquots can also identify discrepancies in the taxonomic annotation selected by alignment and the LCA algorithm.

### tax-aliquots: Towards accurate taxonomic classification and interpretable annotations using homology-based clustering and kAAmer overlap

Combining database curation and unsupervised approaches can improve the accuracy of sequence classification for assembled sequences from meta-omic datasets. Unsupervised approaches have been developed to specifically combat inadequate reference database coverage^38, 39^. However, these approaches tend to rely on classifying highly dissimilar fragments (e.g., separating at the domain level between eukaryotes and prokaryotes) due to genetic overlap among the more taxonomically closely-related sequences. We posit that leveraging large eukaryotic databases, preprocessing the database to reduce problem size and taxonomic overlap, and then training an unsupervised model can improve interpretability of community assessment. Here we leverage clustering tools for a two-stage method of taxonomic assignment, an approach we have named “tax-aliquots: Assigning Lineage to Queries Over Two Steps”. Proteins are first clustered according to their homology, and then hierarchically using the kAAmer (subsequences of amino acids) content of the proteins in the homology-based cluster. The advantages of this method are twofold: we reduce the computational complexity of kAAmer matching^40^, which is an effective tool to distinguish taxonomic groups^41^, and we ensure that assignment is also constrained by sequence alignment.

We tested three distance thresholds for tax-aliquots in the second clustering stage: a permissive, intermediate, and stringent strategy (see Methods). Similar to the percent identity cutoffs used to make decisions about taxonomic level in the Least Common Ancestor (LCA) approach, the distance threshold determines how small the distance between sequences needs to be in order for them to fall into the same cluster. Unlike the LCA approach, all labels are retained in each cluster once they meet the cutoff (Supplementary Figures 13 and 14).

We applied our clustering method to each of the three vignettes discussed above to explore the advantages of using tax-aliquots (Figure 6). In the *Tara* dataset, the tax-aliquots algorithm expands the number of sequences that can be linked to the genus *Phaeocystis* by 47% when a matching species or strain reference is unavailable. 100,879 total *Phaeocystis* sequences were identified by the original BLAST-LCA search containing the full database, while only 11,822 of those sequences were also annotated as *Phaeocystis* using BLAST-LCA with the database containing only free-living *Phaeocystis* references. 6,320 additional *Phaeocystis* sequences fell into tax-aliquots clusters that contain sequences from free-living *Phaeocystis* references. Of these, 2,550 sequences fell into clusters with *Phaeocystis* only (meaning that they were “taxonomically-coherent” for *Phaeocystis*), and the related haptophyte genus *Pavlova* was only the additional member of 84.5% of the remaining clusters. Using the colony-forming database, 5,762 sequences annotated as Phaeocystis using the combined database, but not by the colony-forming database using BLAST-LCA, fell into taxonomically-coherent *Phaeocystis* clusters (a 6.7% increase in annotated sequences).

**Figure 6.**
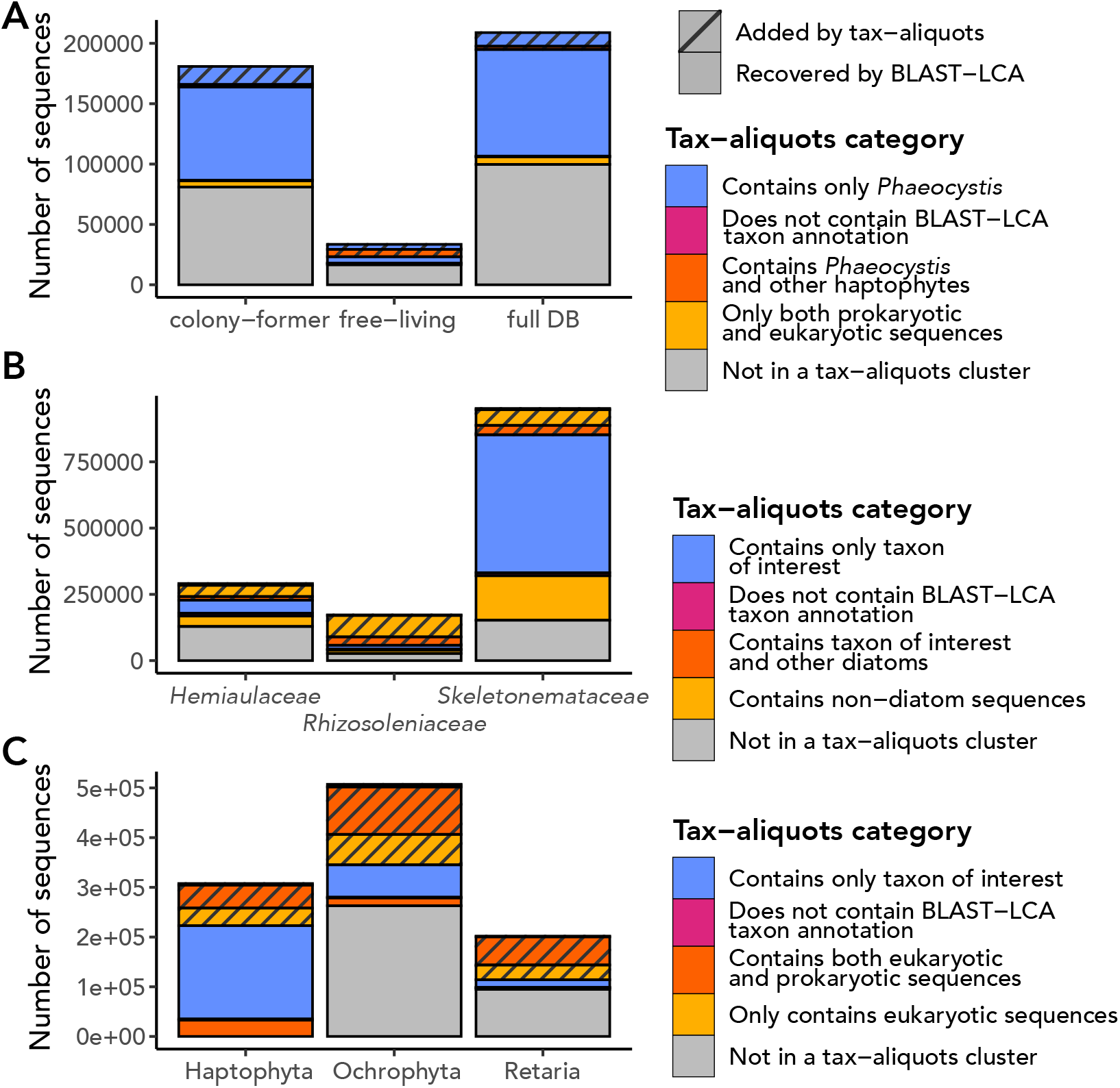
Diagnostic capacity of the tax-aliquots algorithm to refine taxonomic group prediction. All clustering results shown in this figure were found using the intermediate strategy. A: In the Southern Ocean surface water assembly from *Tara* Oceans, the tax-aliquots algorithm identifies sequences that the BLAST-LCA approach does not. These sequences may be *Phaeocystis* or a closely-related haptophyte. Most sequences in coherent *Phaeocystis* clusters not identified by the BLAST-LCA approach were unannotated by BLAST-LCA, not misannotated. B: Comparison of tax-aliquots clusters containing the three families of diatoms considered in vignette 2 (family-level). These clusters overlap and do not constitute an additive community. Cluster sizes demonstrate that a large number of sequences are found in taxonomically-incoherent clusters that include 1) tax-aliquots clusters containing the family of interest and any other member of the class Bacillariophyta, 2) tax-aliquots clusters containing members outside of class Bacillariophyta but within phylum Ochrophyta, or 3) tax-aliquots clusters that contain multiple phyla. Coherent clusters contain only the family of interest. C: Tax-aliquots clusters from the BATS dataset, excluding sequences that were not captured by the first clustering step. Many sequences from the BATS dataset are taxonomically-ambiguous per tax-aliquots, indicative of higher sample diversity and greater divergence from the database sequences, but haptophyte sequences are most likely to be found in coherent clusters. Eukaryote tax-aliquots clusters contain only eukaryotic sequences.

In the Narragansett Bay dataset, the increase in sequences that could be annotated within five families using tax-aliquots, but were unannotated by BLAST-LCA, was a more modest 1.3%. However, the tax-aliquots algorithm offers an explanation for the discrepancy between the metatranscriptomes and the light microscopy in the Narragansett Bay dataset: family *Hemiaulaceae* annotations appear less robust to stringent clustering and are possibly overannotated (Supplementary Figures 13 and 14). Depending on how strictly taxonomy was assigned with tax-aliquots, results had markedly different relative composition of sequences associated with these three main diatom families (Supplementary Figure 11). Further, the tax-aliquots approach represents a powerful technique to expand understanding of unknown sequences in collaboration with alignment-based searches. As an example, of the 2,942,183 sequences found in clusters containing references from one of the three diatom families, 1,046,573 were unannotated at the family level by BLAST-LCA, and 69,275 of those sequences were unannotated at the domain level. Among the sequences unannotated at least at the family level, 526,897 fell into permissive tax-aliquots clusters, including 121,748 that fell into clusters containing a single family.

The tax-aliquots method reveals that even after radiolarians were added to the database, the reference database is insufficient to accurately annotate taxonomy in diverse samples. Using the annotation strategies mentioned here, the BATS dataset continued to have a low number of successful taxonomic annotations for sequences that co-clustered with Retaria. Tax-aliquots revealed that few putative radiolarian sequences correspond to taxonomically unambiguous clusters, even though a similar number of putative radiolarian sequences fell into Retaria taxonomic clusters (Supplementary Figure 12). Retaria also has far fewer taxonomically-coherent clusters than dominant phyla from the same sample (e.g., Haptophyta, Supplementary Figures 13 and 14). The majority of ambiguous radiolarian sequences fell into clusters that also contained dinoflagellates and foraminifera (Supplementary Figure 12), a problem we also noticed when adjusting the database and using the DIAMOND/BLAST and LCA approach. This observation is likely due to database incompleteness and high taxonomic overlap between adjacent taxa (Supplementary Figure 12). Sequences that were affected by the database change were more likely to be ambiguously labeled by tax-aliquots. For example, of 3,025 sequences that were annotated as Ochrophyta before but not after adding in Retaria, only 3% were in coherent tax-aliquots clusters (93), as compared to 10.7% overall (35,694 of 333,210 Ochrophyta proteins). Because tax-aliquots reflects taxonomic ambiguity in the outcome of clustering, misinterpretation is less likely.

## Discussion

The growth of databases and the development of complementary computational analysis approaches has enabled taxonomic predictions for community assessment in meta-omics. The overall size of available databases has expanded dramatically since the first environmental metagenome, fueled by the growing availability of genomes and new sequencing technology that can be deployed straight from the lab (*e.g.*, Nanopore sequencing^42–44^), and the curation of resources from transcriptomes^19, 27, 29, 36, 45–47^ and metagenome-assembled genomes^9^ for eukaryotes^10–12, 48^. Improved sampling and sequence curation have accelerated the development of annotation approaches that can accurately assess the whole community, leveraging databases of predicted genes or full contigs for taxonomic classification beyond the use of marker gene alignments.

Database curation plays a critical role in how sequences are taxonomically annotated, and how taxonomic identity is linked to functional role. Researchers need to be cognizant that all database mapping is selective: bias inherently exists in all taxonomic mapping, as only a selection of organisms have been isolated, subsequently sequenced, and added to reference databases. Because microeukaryotes have high average genetic differentiation^49^, much of our ability to annotate diversity hinges on tradeoffs inherent to building appropriate databases. The annotation of Radiolaria in the BATS transect was only made possible by the addition of Radiolarian references present only in the EukProt and EukZoo databases^37, 47^, as no Radiolaria are present in the MMETSP^29^. Further, after applying the tax-aliquots approach, it became clear that database completeness still limits precise annotation. Hence, it is likely that other understudied microeukaryotic lineages are present in the North Atlantic dataset and other meta-omic datasets. Further, these missing references can completely change the interpretation of relative community composition, even to the point of reshaping predicted taxonomic affiliations outside of the missing group that was added.

However, database expansion is not always the solution. We found that more than half of sequences within major phyla (e.g. Bacillariophyta) lack non-self hits to another sequence of the same family (Table 1; Supplementary Figure 6). Because non-self, same-family hits appeared to be limited to a maximum value regardless of the number of available family-level relatives in the database (Supplementary Figure 6), this observation is unlikely to be solely a consequence of database incompleteness. In some cases, the sequences lacking family overlap might be spurious, and in other cases sequences may constitute valuable variability that could enable understanding of population dynamics in protists^50, 51^. The importance of database completeness and expansion is made clear by the effect of the presence or absence of particular species in the *Phaeocystis* database mapped against the *Tara* Oceans samples. The addition of genomes and transcriptomes at genus resolution did not necessarily increase our ability to identify a different species from that genus using typical annotation approaches. Further, when it comes to protein matching, percentage identity within a high-scoring alignment is frequently an unreliable indicator of phylogenetic relatedness.

**Table 1.**
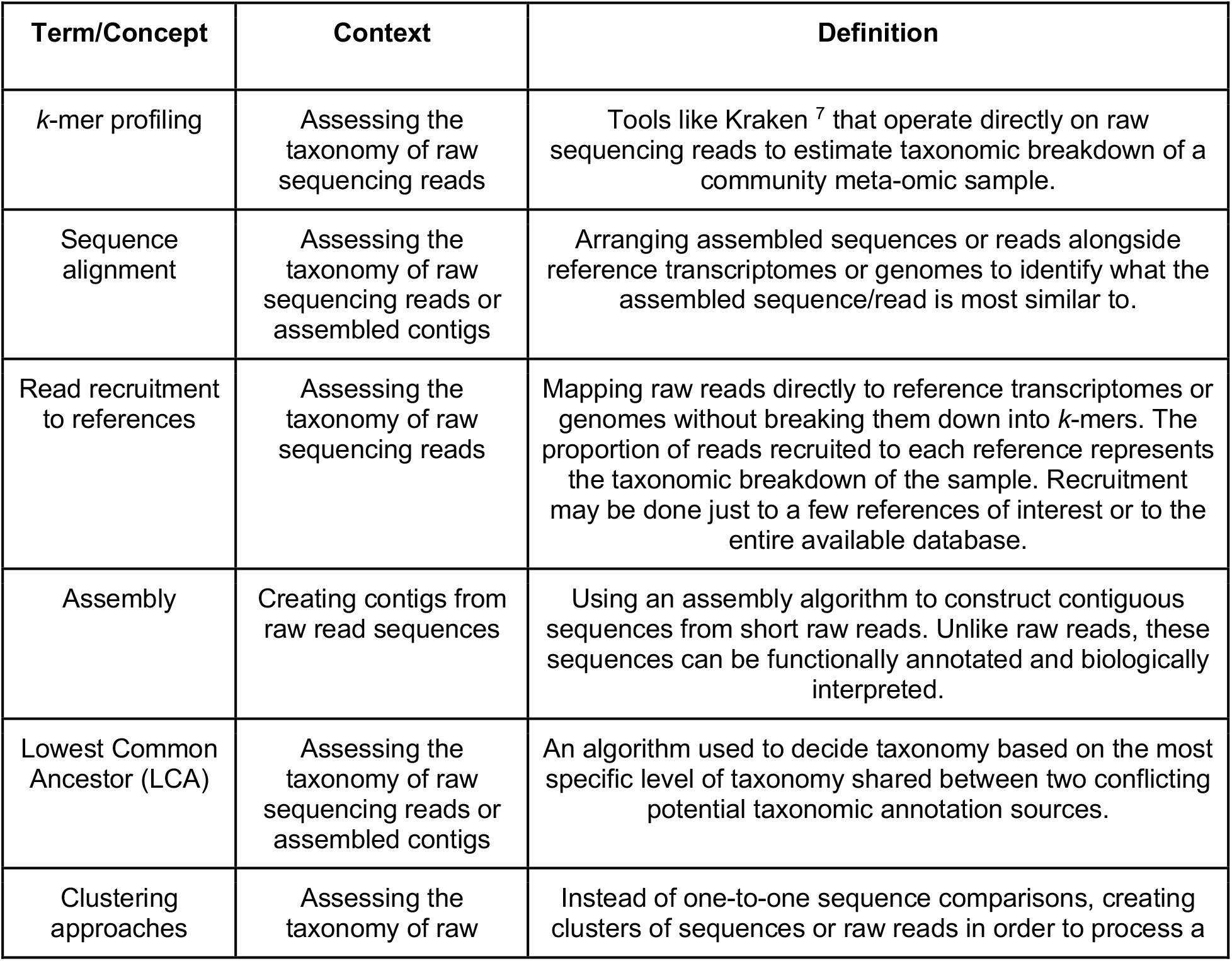
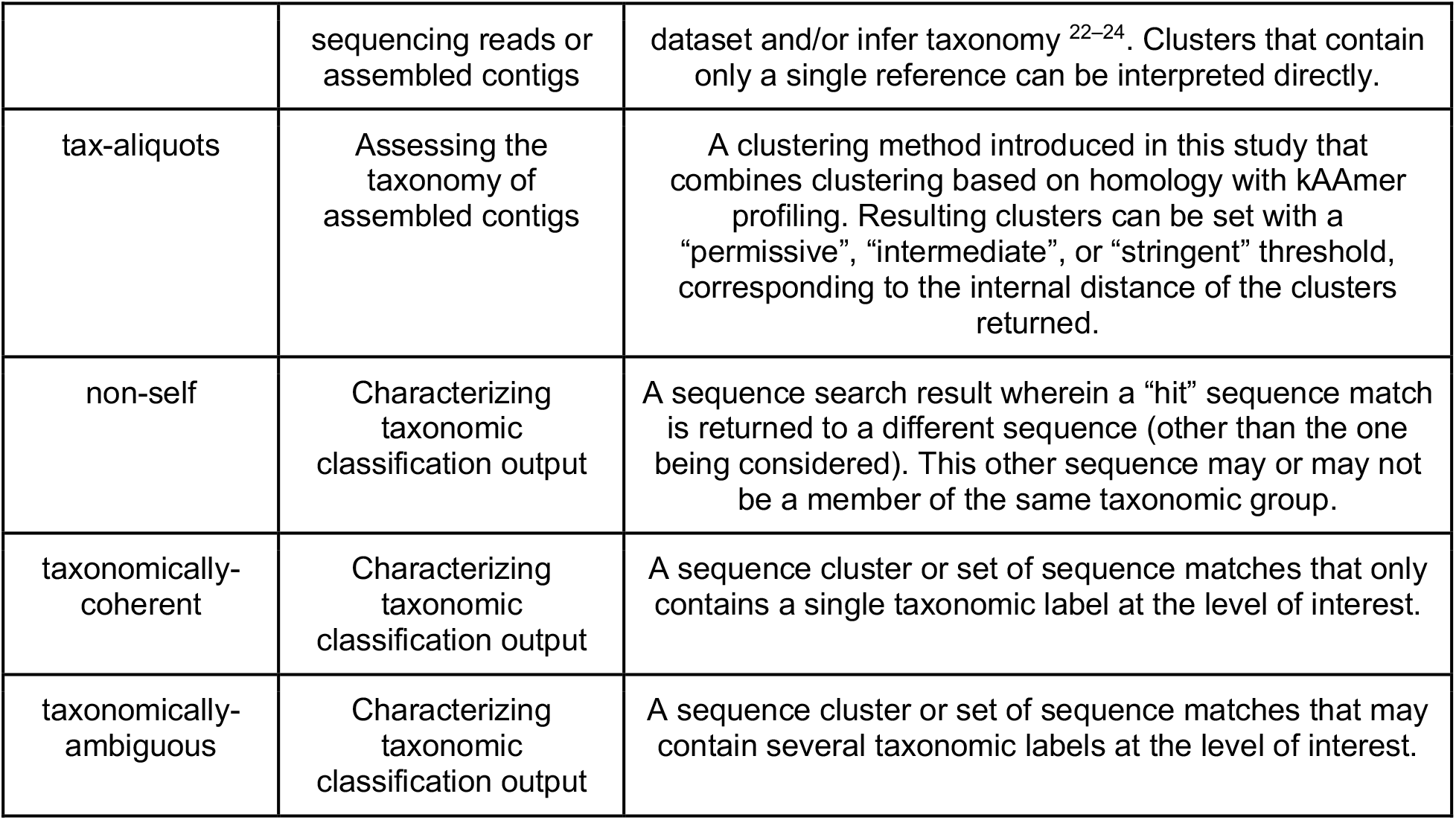
Summary of terms used in the paper to describe methods to annotate meta-omic sequencing datasets.

The future of taxonomic classification in multi-omic studies must balance the growing availability of database sequences with computational approaches to decipher their origins. Approaches that have been available since the early days of metagenomics, like Naïve Bayes classification^52, 53^, deep learning, and topic modeling have become less popular in recent literature in favor of more direct comparisons to databases, which are more interpretable but also minimally predictive^54–56^. The unbalanced (with respect to taxonomic distribution across the tree) number of available references for different phyla and orders necessitates pre-processing and careful model training (Supplementary Figure 2). Training models or selecting thresholds using a phylogeny-aware approach also takes into account the patterns in sequence overlap that differentiate microorganisms (*e.g.*, what defines distinct species at the sequence-level for one family may be different for another family). In more remote environments such as the deep sea, in which a smaller proportion of sequences are expected to have complete database counterparts, using a generative and flexible approach such as topic modeling or global hierarchical clustering (instead of a homology search) may be warranted.

Accurate taxonomic annotation of environmental sequences is a dynamic problem, which has evolved with both algorithms and the increasing size of databases. Here, we propose a hybrid approach that is meant to complement alignment and LCA-based approaches and to identify gaps in annotation accuracy. “Tax-aliquots” combines alignment-based estimation using large databases with computed kAAmer similarity over the entire sequence. Our hybrid approach is also equipped to maximize future improvement as reference databases grow. Using an unsupervised method and a clustering approach reduces bias associated with particularly rare taxonomic groups for which only a single database representative might be available. Because all sequence matches are treated equally, multiple repeated hits are not weighted more heavily, allowing for the identification of annotation challenges. Taken together, our vignettes and the output of tax-aliquots illustrate the importance of critically evaluating the completeness and composition of the database selected. Tax-aliquots and other algorithms are essential tools to increase annotation accuracy and avoid the pitfalls of incomplete databases. Tax-aliquots provides an approach to target taxa that require expanded database coverage to be identified. We encourage applying the open-source tax-aliquots workflow to challenging datasets with low rates of taxonomic annotation, and to databases themselves to identify indistinguishable overlaps between groups and make taxonomic assignment of diverse microbial eukaryotes more interpretable. Critical reassessment of datasets and evaluation of methods is a vital step towards linking taxonomic variability to functional potential in *in situ* communities of ecologically-essential protists.

## Code availability

Code for running the tax-aliquots clustering can be accessed at https://github.com/akrinos/tax-aliquots

## Acknowledgements

This material is based upon work supported by the U.S. Department of Energy, Office of Science, Office of Advanced Scientific Computing Research, Department of Energy Computational Science Graduate Fellowship under Award Number DE-SC0020347 (A.I.K.). The work conducted by the US Department of Energy Joint Genome Institute (https://ror.org/04xm1d337), a DOE Office of Science user facility, is supported by the Office of Science of the US Department of Energy, operated under contract no. DE-AC02-05CH11231 (F.S., A.I.K.). M.M.B. is supported by a Simons Foundation Postdoctoral Fellowship in Marine Microbiology (award 874439). H.A. is supported by a Simons Foundation Early Career Investigator in Aquatic Microbial Ecology and Evolution Award (award 931886).

## Online Methods

In order to evaluate and select a sequence identity cutoff for use in taxonomic classification, we performed a bidirectional DIAMOND search^57^ of the MMETSP database using the blastp algorithm^58^. We used a cutoff of hits with bitscore of at least 50, and processed hits according to their percentage identity. We removed self-hits to the same sequence, and then recorded the percentage of sequences within each taxonomic family that had (a) hits to other sequences in the same taxonomic family and (b) hits to other sequences in different taxonomic families using eight different percentage identity cutoffs (30,40,50,60,65,70,80, and 90). We compared each of these percentages to the total number of transcriptomes associated with each family within the MMETSP. The results from this bidirectional search were used for the diatom family best hits displayed in Figure 1D and for the diatom family mean percentage identity results in Figure 3B. A similar bidirectional search which also included additional Radiolarian references was used to generate Supplementary Figure 2E, and the same bidirectional search among the *Phaeocystis* references above was used to generate Supplementary Figure 2F. We tested the uniformity of the counts of each diatom family in the MMETSP using the Anderson-Darling test against the uniform distribution generated with a count bound of zero to 10 greater than the maximum observed per-family count using the goftest package (version 1.2-3) in R^59^.

### Genus Scale: Tara Oceans metagenomes

Metagenomic samples from the global ocean were retrieved from the *Tara* Oceans project^60^. Assemblies were previously generated in Alexander et al. (2021)^12^, with input sequencing reads grouped by ocean basin, depth, and size fraction; in brief, assemblies were generated by the MEGAHIT assembler^61^ after trimming with the Trimmomatic software^62^. Protein prediction was performed with Prodigal^39, 63^. The taxonomic identity of predicted proteins was obtained using EUKulele v2.0.3^19^, first using a combined database containing the MMETSP^27, 29, 36^, MarRef^64^, and additional *Phaeocystis* references, including the genome resources for *Phaeocystis antarctica* and *Phaeocystis globosa*^65, 66^ available from the IMG/M (Integrated Microbial Genomes & Microbiomes) database (Phaant1 and Phaglo1, respectively), *Phaeocystis cordata*, *Phaeocystis jahnii*, and *Phaeocystis globosa* transcriptome resources^67–69^, and a *Phaeocystis pouchetii* transcriptome (Mars Brisbin et al. *in prep*). The contigs associated with the proteins identified to the genus *Phaeocystis* were quantified against the raw reads using the CoverM software in contig mode (v0.6.2; https://github.com/wwood/CoverM; coverm contig ––min-covered-fraction 0).

Subsequently, separate EUKulele databases were created that contained the MMETSP^27, 29, 36^ with all genus *Phaeocystis* references removed, the MarRef^64^ database, and one of the ten distinct *Phaeocystis* genome or transcriptome references, inclusive of species *Phaeocystis antarctica*, *Phaeocystis globosa*, *Phaeocystis pouchetii*, *Phaeocystis jahnii*, *Phaeocystis cordata*, and *Phaeocystis rex*. A third set of EUKulele databases was created which contained the MMETSP^27, 29, 36^ with all genus Phaeocystis references removed, the MarRef^64^ database, and all of either the colony-forming *Phaeocystis* species or the free-living *Phaeocystis* species (*Phaeocystis cordata*, *Phaeocystis jahnii*, and *Phaeocystis rex*). Each *Tara* Oceans assembly was annotated with each of these databases.

A phylogenetic tree for the *Phaeocystis* references was constructed by conducting orthologous group clustering against all *Phaeocystis* references, a selection of *Emiliania huxleyi* transcriptome assemblies from the MMETSP (MMETSP0994, MMETSP0995, MMETSP0996, MMETSP0997, MMETSP1006, MMETSP1007, MMETSP1008, MMETSP1009, MMETSP1150, MMETSP1151, MMETSP1152, MMETSP1153, MMETSP1154, MMETSP1156, MMETSP1157), *Gephyrocapsa oceanica* transcriptome assemblies from the MMETSP (MMETSP1363, MMETSP1364, MMETSP1365, MMETSP1366), *Isochrysis galbana* transcriptome assemblies from the MMETSP (MMETSP0943, MMETSP00595), and three reference genomes from the JGI’s IMG/M (Integrated Microbial Genomes & Microbiomes) database^65, 66^ – *Chrysochromulina tobinii* (Chrsp), *Oxytricha trifallax* (Oxytri1), and *Guinardia theta* (Guith1). Orthologous groups were created from proteins from all references using OrthoFinder (v2.5.4)^70^, and orthologous groups containing a single protein from all of the *Phaeocystis* references were used to create an alignment and phylogenetic tree. This amounted to 40 total single-copy genes shared across references which were used to build the alignment. The MAFFT tool was used for multiple sequence alignment of each of the concatenated lists of single-copy genes (one file per gene containing all gene versions across organisms in the alignment; version 7.508), followed by the removal of possible spurious sequences using trimAl^71^ (version 1.4.rev15), and then a secondary multiple sequence alignment using Clustal-Omega^72^. Sequences in the alignment were adjusted to standardize their trimmed lengths, and the subsequent alignments were concatenated and trimmed once more with trimAl. FastTree (version 2.1.11) was used to build the phylogenetic tree^73^.

### Family Scale: metatranscriptomes from Narragansett Bay

The metatranscriptome assembly and annotation process for the metatranscriptomic samples from Narragansett Bay is described in full in Krinos et al. (2023)^35^. In brief, raw reads were trimmed and quality-assessed, and then assembled in parallel using the eukrhythmic pipeline^35^. Taxonomic annotations were assigned using the EUKulele tool^74^ using a combined database containing the MMETSP and MarRef2 sequences^29^.

### Phylum Scale: metatranscriptomes from a transect between WHOI and BATS

Samples from the transect between Woods Hole Oceanographic Institution (WHOI) and the Bermuda Atlantic Time Series (BATS) stations were assembled and post-processed as described in Cohen et al. (2023; *in prep*). EUKulele^74^ was used for the BLAST-LCA search against these sequences, first using the MarRef and MMETSP database^29^ and then adding all radiolarian references available in the EukProt and EukZoo databases^37, 47^. These organisms included *Sticholonche zanclea* (EP00491), *Amphilonche elongata* (EP00492), *Phyllostaurus siculus* (EP00493), *Astrolonche serrata* (EP00494), *Collozoum sp. 1* RS2012 (EP00495), *Lithomelissa setosa* (EP00496), and *Spongosphaera streptacantha* (EP00497).

### Hybrid partially-supervised clustering workflow

A very permissive protein clustering is performed using DIAMOND DeepClust^23^, followed by taxonomic profiling using hierarchical clustering on a matrix formed in parallel by calculating kAAmer overlap between sequences present in the cluster. This enables exact kAAmer overlap to be computed efficiently, and does not taxonomically annotate sequences for which an alignment is based on sequence coverage of <20-50% of the protein. Unlike other LCA-based approaches where ancestry is computed using the aligned fragment, this method profiles the short kAAmers over the entire length of the proteins which were originally clustered together on the basis of a short and potentially low sequence similarity alignment. This allows sequences with promising homology, even with low percentage identity, to be clustered based on consistency in sequence content over the entire protein length.

We ran DIAMOND DeepClust^23^ against the predicted proteins from the MMETSP and MarRef2 databases^29^ using a 50% coverage threshold for the shorter sequence in the alignment and no minimum percentage identity. First, kAAmers were identified in parallel separately for each cluster. We used the pyahocorasick package, which implements the Aho-Corasick algorithm for efficient string matching^75, 76^. After counting all kAAmers of length 4 using this approach and the Automaton utility from pyahocorasick, we computed similarity between each sequence in the protein cluster according to the formula:

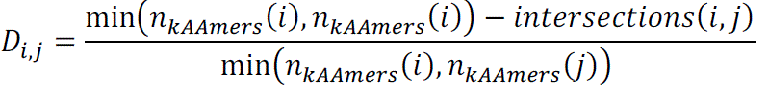

Where *intersections*(*i*, *j*) is the number of intersecting kAAmers between proteins sequences *i* and *j* and min(*n_kAAmers_*(*i*), *j_kAAmers_*(*i*)), is the minimum number of kAAmers found in each of the two protein sequences, which is used to scale the raw number of intersections. These distance numbers were used for the downstream hierarchical clustering steps, which were conducted using the fcluster function from SciPy^77^.

We linked original sequences from the database to revised taxonomic annotations according to the taxonomic coherence of the cluster to which it was assigned using the two-part algorithm. We created a new taxonomy string dictionary which takes into account the taxonomic ambiguity of sequences according to their kAAmer overlap. Then, we applied this new taxonomy string to best hits from the Narragansett Bay (Family Scale) and BATS (Phylum Scale) datasets which were originally annotated using the MarRef2 and MMETSP database proteins and using a kAAmer length of 3. The stringent approach used a distance threshold of 0.2, the intermediate a threshold of 0.5, and the permissive approach used a distance threshold of 0.8.

Figures were generated in R (version 4.1) and in Python (version 3.10.1) using the ggplot2 software, ggridges package, ggUpSeT package, ggmaps package, and ggalluvial package^78–83^.

