## Supplementary Information for "Missing microbial eukaryotes and misleading meta-omic conclusions"

**Supplemental Material for “Missing microbial eukaryotes and misleading meta-omic conclusions”**

**Genus Scale**: Transcriptomic and genomic references from closely-related species do not imply genus-level resolution in *Phaeocystis*

Some microbial eukaryotes have relatively small evolutionary distance, yet distinctive geographic distributions and morphologies. One such eukaryotic group is the Haptophyte genus *Phaeocystis*. *Phaeocystis globosa* is the most globally-ubiquitous species of *Phaeocystis* and shares genes with many different species of *Phaeocystis*. *Phaeocystis antarctica* is most phylogenetically similar to fellow colony-forming species *Phaeocystis pouchetii* [31], yet the two are morphologically distinct and found at opposite poles [32]. In contrast, *Phaeocystis cordata* and *Phaeocystis jahnii* are both found at mid-latitudes, do not form colonies, and are more phylogenetically similar to one another than they are to any other *Phaeocystis* species [33,34].

In *Tara* Oceans samples from the Mediterranean Sea and the Southern Ocean, the sequence annotation rate in the genus *Phaeocystis* varied depending on the number and ecotype of *Phaeocystis* reference genomes and transcriptomes used. In Mediterranean Sea samples where free-living *Phaeocystis* species tend to be dominant [32–34], databases which contained one or more free-living *Phaeocystis* references resulted in a greater number of sequences annotated at the genus level to be *Phaeocystis*. In the surface Southern Ocean samples where colony-forming Phaeocystis antarctica is typically dominant, an additional 67% of sequences could be annotated as genus *Phaeocystis* using a database containing only the colony-forming ecotypes as compared to one containing only the free-living ecotypes (Figure 3B). This amounted to 86,097 sequences (79.0% of total *Phaeocystis*-identified sequences) using the colony-former database, as compared to 12,307 sequences (11.3% of total *Phaeocystis*-identified sequences) using the free-living database. In the Mediterranean Sea surface small size fraction, 58.4% of sequences annotated to be *Phaeocystis* when using a database containing all available *Phaeocystis* references were also annotated using the free-living *Phaeocystis* database (52,312 sequences; Figure 3C), while only 39.9% of all sequences were annotated using the colony-former database. The free-living database outperformed the colony-former database in both the small- and large- size fraction samples (Figure 3C). This may imply that size fraction is a poor indicator of the extent of colonization, given that these organisms are not likely to form colonies and that colony forms are rarely observed in the Mediterranean Sea [84]. The overall number of sequences was much lower in the Mediterranean Sea than in the Southern Ocean and the difference between databases was less pronounced (152 vs. 289 sequences annotated using the colony-forming and free-living databases, respectively).

One of the aims of meta-omic analyses is to evaluate changes in taxa across global ocean gradients, yet it is exceptionally difficult to reliably distinguish between species of the genus *Phaeocystis* (Figure 3) using conventional methods of taxonomic assignment [31,68]. Studies which leverage large-scale global datasets to infer taxonomic distribution from metagenomes or metatranscriptomes [15,85–87] are limited by the target level of taxonomic resolution and the quality of genomic and transcriptomic resources used to create the database. *Phaeocystis* is an ecologically-relevant example for the importance of database expansion, in particular when transcriptomic references without an exhaustive gene set are used.

**Family Scale**: Different references with uneven results in tracking diatom (class *Bacillariophyta*) blooms in Narragansett Bay

A major goal and utility of meta-omic surveys is to increase sample throughput and accurately identify organisms that might not have been differentiable or countable in microscopic samples. Pairing meta-omic data to microscopic counts, especially for numerically abundant groups, can help to target database expansion efforts. Here we provide the example of Narragansett Bay, the site of a Long-Term Plankton Time Series (University of Rhode Island Graduate School of Oceanography Plankton Time Series; https://web.uri.edu/gso/research/plankton/data/) where a metatranscriptomic survey was conducted and published in 2015 [26]. These metatranscriptomes were originally taxonomically annotated using the raw read recruitment approach to the Marine Microbial Eukaryote Transcriptome Sequencing Project (MMETSP) database [26,27,29,36], and were later re-analyzed using a metatranscriptome assembly approach [88].

The dominant diatom cataloged in the microscopic count data was not consistently detected by metatranscriptome-derived community expression (class *Bacillariophyta*; [26]. On two dates in the time series, the two dominant diatoms in the microscope data were *Dactyliosolen fragilissimus* (family *Rhizosoleniaceae*) and *Cerataulina pelagica* (family *Hemiaulacaeae*; University of Rhode Island Graduate School of Oceanography Plankton Time Series; https://web.uri.edu/gso/research/plankton/data/). The diatom *Cerataulina pelagica* is absent from the MMETSP database, hence it was not accurately identified in the metatranscriptomic data at any level except at the family level (*Hemiaulaceae*). This is problematic because family *Hemiaulaceae* is extraordinarily diverse, found globally in water temperatures from –2 to 30 degrees Celsius, with different members having different ecological roles (*Encyclopedia of Life*; https://eol.org/pages/3473). *Dactyliosolen fragilissimus* is present in the MMETSP database (Figure 2), but a very low proportion (<1%) of species-level annotations were retrieved. In addition to *D. fragilissimus,* three of its relatives at the family level *(Rhizosoleniaceae*) are also included in the MMETSP database (*Rhizosolenia setigera*, *Proboscia inermis*, and *Proboscia alata*; Figure 2), yet less than 5% of reads were attributed to this family at the peak abundance of *D. fragilissimus* in the metatranscriptomic samples (Figure 5A). Despite the availability of a laboratory-grown *Dactyliosolen* reference in the database, taxonomic resolution is inadequate in the meta-omic record of the sample [27,29]. The outcomes of low database taxonomic resolution were incongruent between taxa: though both missing taxa of *Hemiaulaceae* and *Rhizosoleniaceae* had a member of the same family available in the database, only *Hemiaulaceae* yielded annotations at the expected taxonomic resolution. Critically, this implies that taxonomic coverage alone often does not lead to accurate taxonomic labels.

Parsing the alignment results provided further evidence that *Rhizosoleniaceae* may be inadequately reported. While 318,420 (4.2%) ORFs in the Narragansett Bay samples using the combined assembly had a hit with bitscore of more than 50 and percentage identity of more than 65% to *Rhizosoleniaceae*, only a fraction of those ORFs (30,612; 0.4% of total) were assigned a *Rhizosoleniaceae* annotation via the LCA algorithm (total ORFs: 7,624,081). Together with the evidence that a maximum of 6,139,000 cells were estimated to belong to *Rhizosoleniaceae* per liter of water in the microscopic count data, this observation prompts further exploration of whether *Rhizosoleniaceae* was under-annotated computationally rather than under-sequenced in the metatranscriptomes.

A significant decision for ‘omics researchers is whether metatranscriptome annotation is applied to ORFs using the blastp algorithm or full contigs using the blastx algorithm: the blastx algorithm will translate the nucleotide sequence into all six possible reading frames and return the best hits to any of those reading frames, resulting in a single taxonomic annotation per contig [57,58]. By contrast, using TransDecoder (https://github.com/TransDecoder/TransDecoder/wiki) or similar to identify ORFs prior to annotation may result in different taxonomic annotations on each contig, and requires reads to be split and assigned to ORFs from full contig mapping. This becomes complex with respect to considering ORF length and whether all ORFs from the contig have been assigned annotations. In the Narragansett Bay dataset, we found that 31,554 (0.25% of total annotated contigs) contigs contained ORFs with taxonomic annotations that conflicted at the family-level to the best annotation identified on the contig. A far more likely outcome was that at least one coding sequence did not have any family-level annotation despite the full contig being annotated at the family-level (4,646,475 contigs; 35.6% of total). The reverse scenario (a coding sequence being annotated at the family-level whereas the full contig was not, likely due to a conflict) was rarer (57,907 contigs; 0.44% of total). 5,141,182 contigs (39.3%) were lineage-conflicted at the family-level, i.e., having a classification at a higher level of taxonomy than the family-level, but having conflicts in both coding sequence and contig. Some families are more significantly impacted than others: among frequently-annotated families, 1.35% of contigs identified as family *Hemiaulaceae* (1,819 of 134,950) and 1.32% of the *Strombidinopsidae* (1,218 of 92,403) via the blastx algorithm had lineage conflicts among ORFs, whereas only 0.08% (579 of 702,145) of the family *Skeletonemataceae* and 0.07% (116 of 164,309) of the *Leptocylindraceae* did. In sum, assigning the consensus family annotation of the contig to all coding sequences present on the contig only generated a conflict 0.24% of the time, yet could result in new family-level taxonomic annotations for 35.5% of ORFs.

Cutoff choice affects the level of taxonomy assigned and hence the ability of researchers to interpret community composition [89]. Phylogenetic distance has been well-established for many metabarcoding studies. In metagenomics and metatranscriptomics, the boundaries are poorly defined, as individual genes may vary in divergence. A more lenient cutoff will increase conflicts, while a more stringent cutoff may prevent crucial links from being made when a given sequence is absent for the target organism but present in its relatives. We find that a percentage identity threshold of 80% optimizes family-level taxonomic annotation of microbial eukaryotes using a transcriptome reference database. Above this threshold, the proportion of sequences with a non-self hit within the same taxonomic family begins to fall, reducing the likelihood that another sequence will be available for annotation when one from the target taxon is unavailable. Below this threshold, conflicts of non-self hits to different taxonomic families rise above 5% of sequences. Approximately half of all sequences in a taxonomic family appear to be reference-specific and will not be found unless that specific sequence or a relatively rare related sequence is added to the database. Additionally, we found that having at least 10 transcriptomes from a taxonomic family of interest within eukaryotes resulted in having an average of more than half of sequences having an accepted hit to another sequence in the database of the same taxonomic family. Beyond this number of references, the marginal benefit of adding additional references becomes saturated (Supplementary Figure 6).

Underrepresented families in the MMETSP database were also likely to be misannotated in the bidirectional DIAMOND BLAST search. This was due to a) normalized number of hits to family-annotated references in the database, b) maximum bitscore of a hit to a family, or c) the LCA algorithm applied to all database hits. For example, for family *Hemiaulaceae*, which contains a single transcriptome reference in the MMETSP database, only 69 proteins had a best non-self bitscore hit to another *Hemiaulaceae* sequence with a bitscore of 50 and percentage identity of more than 80%, while 290 sequences had a best non-self bitscore hit to a family that is not *Hemiaulaceae*, of 10,175 total protein sequences. Despite this, *Hemiaulaceae* was the second most-annotated taxonomic family for proteins extracted from the Narragansett Bay dataset, suggesting that these poorly-performing sequences with respect to bidirectional database search can still be environmentally-relevant.

**Phylum Scale**: Missing references impair community assessment along a transect from BATS to WHOI

Approximately 1,021,229 (8.6%) of ORFs were annotated at the domain–but not the phylum–level (lineage-conflicted; either due to percentage identity cutoff or collision between potential annotations). Interestingly, a greater proportion of these phylum-level lineage-conflicted ORFs (95.8%) were assigned a functional annotation using the eggNOG-mapper software as compared to the overall proportion of sequences with a functional annotation (45.8%). This suggests that more conserved proteins will be left out of lineage-specific analysis because they tend to be taxonomically-ambiguous. The analysis along the BATS transect was originally annotated using a database of the MarRef sequences and the MMETSP [29,90]. However, as the MMETSP database does not contain Radiolarians (phylum Retaria; [29] annotating the predicted proteins with the MMETSP led to subsequently identified Radiolarian proteins to be unannotated (n=42,736) or annotated as a different phylum (n=46,283) (Figure 4C). Adding Radiolarian sequences to clustering with DIAMOND DeepClust [23] not only reduced the number of singleton clusters but also reduced the total number of clusters containing sequences that were putative Radiolarians in the EUKulele search (9,548 vs. 7,127 sequences, respectively; Supplementary Figure 7). Just 93 of those clusters contained a MarRef/MMETSP database sequence before and after adding Radiolarians to the clustering, indicating that the majority of reduction (2,328 clusters) was attributable to singleton or small in situ-only clusters being added to larger clusters of sequences (Supplementary Figure 8).

Because functional interpretation is the downstream goal of many omics analyses, it is necessary to predict proteins from metagenomic and metatranscriptomic sequences. However, using only those sequences which contain predicted proteins results in an incomplete picture of community composition. Predicted eukaryotic proteins with known functional annotation (i.e. KEGG/Pfam) were more likely to have lineage-conflicts. This observation implies that shortcomings in taxonomic specificity, whether it be due to indistinguishable overlap between phylogenetic relatives, database inadequacy, or challenges in annotation methodology, disproportionately impacts the functional interpretation that is the goal of many meta-omic studies. Proteins can also be highly conserved and have a degree of sequence content overlap impossible to disambiguate between taxa. Much greater proportion of the original library can be assigned taxonomy when annotation is contig-based. However, this approach lends less directly to the coding sequence-based functional analysis which is often the next step in a multi-omic workflow.

*Hybrid partially-supervised clustering workflow*

Using an alignment approach alone, taxonomic assignment may be possible on the basis of a short fraction of the sequence having high fidelity similarity to a database sequence without the remainder of the fragment being accordingly similar. Using our approach, taxonomy is decided using an exact kAAmer inventory over the whole sequence after alignment, after clustering is used to group sequences. This retains the utility of sequence similarity searching and alignment, but uses the entire sequence to decide taxonomic origin. An identical workflow can be used to identify the taxonomic origin of nucleotide sequences from contigs, which avoids the step of protein prediction that excludes many original contigs. The same two-stage clustering algorithm would efficiently cluster nucleotide sequences and decide within-cluster taxonomy using overlap in longer k-mers.

2,942,183 of 7,624,296 total protein sequences clustered with a database sequence in diatom family *Skeletonemataceae*, *Thalassiosiraceae*, *Hemiaulaceae*, and/or *Rhizosoleniaceae* using DeepClust.

### Supplementary Figures

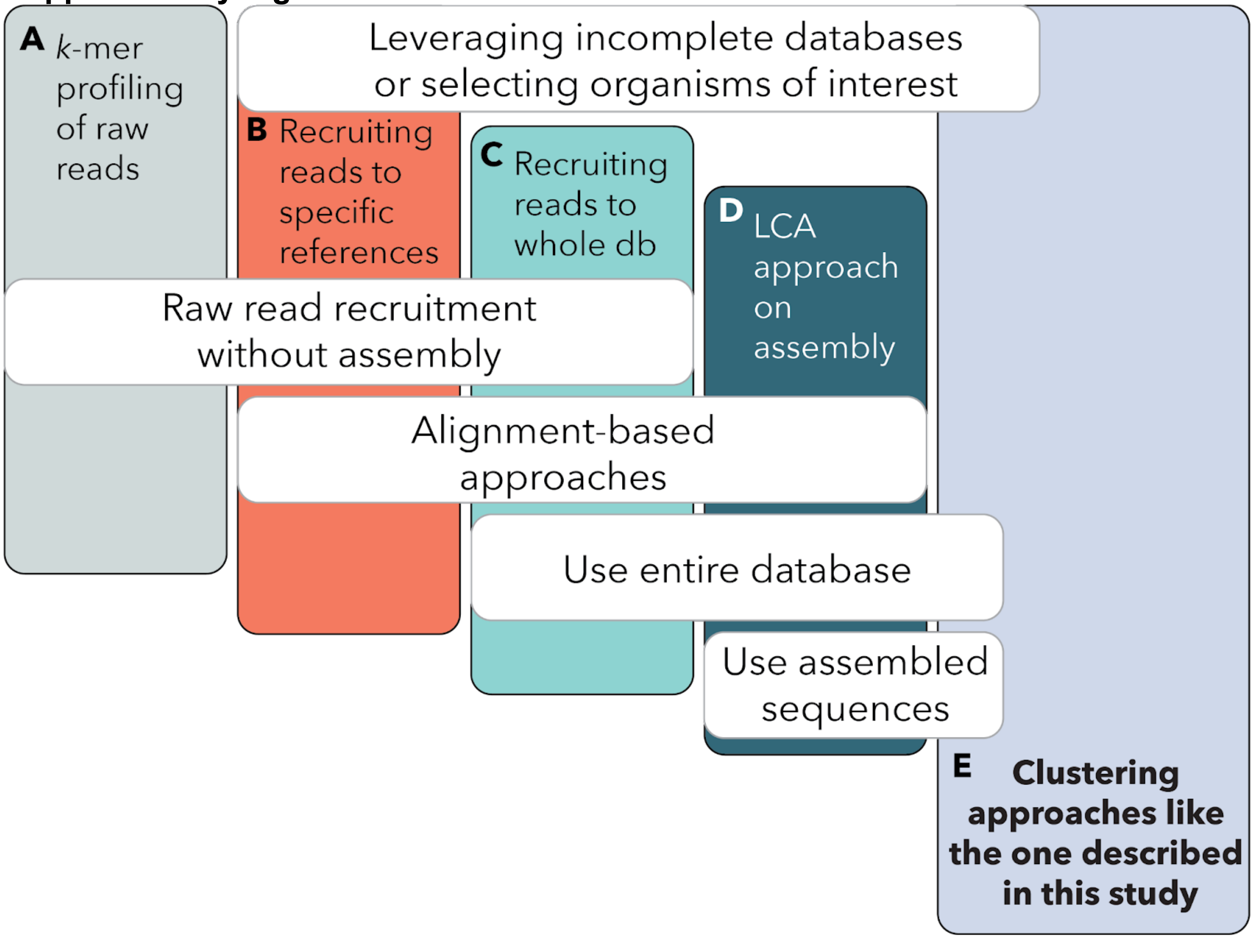


Supplementary Figure 1. Approaches for taxonomic annotation of multi-omic datasets. A: k-mer profiling that attempts to identify taxonomy from raw reads, B: raw read recruitment (without *de novo* assembly) to assembled references from only a selection of organisms believed to be present in the sample or of interest to the research group, C: raw read recruitment (without *de novo* assembly) to a reference database, D: assembly and annotation using a lowest common ancestor approach to a reference dataset, E: our novel hybrid and partially-supervised approach of assessing taxonomy and database limitations in which homology-based clustering of proteins is combined with taxonomic profiling based on subsequence overlap.

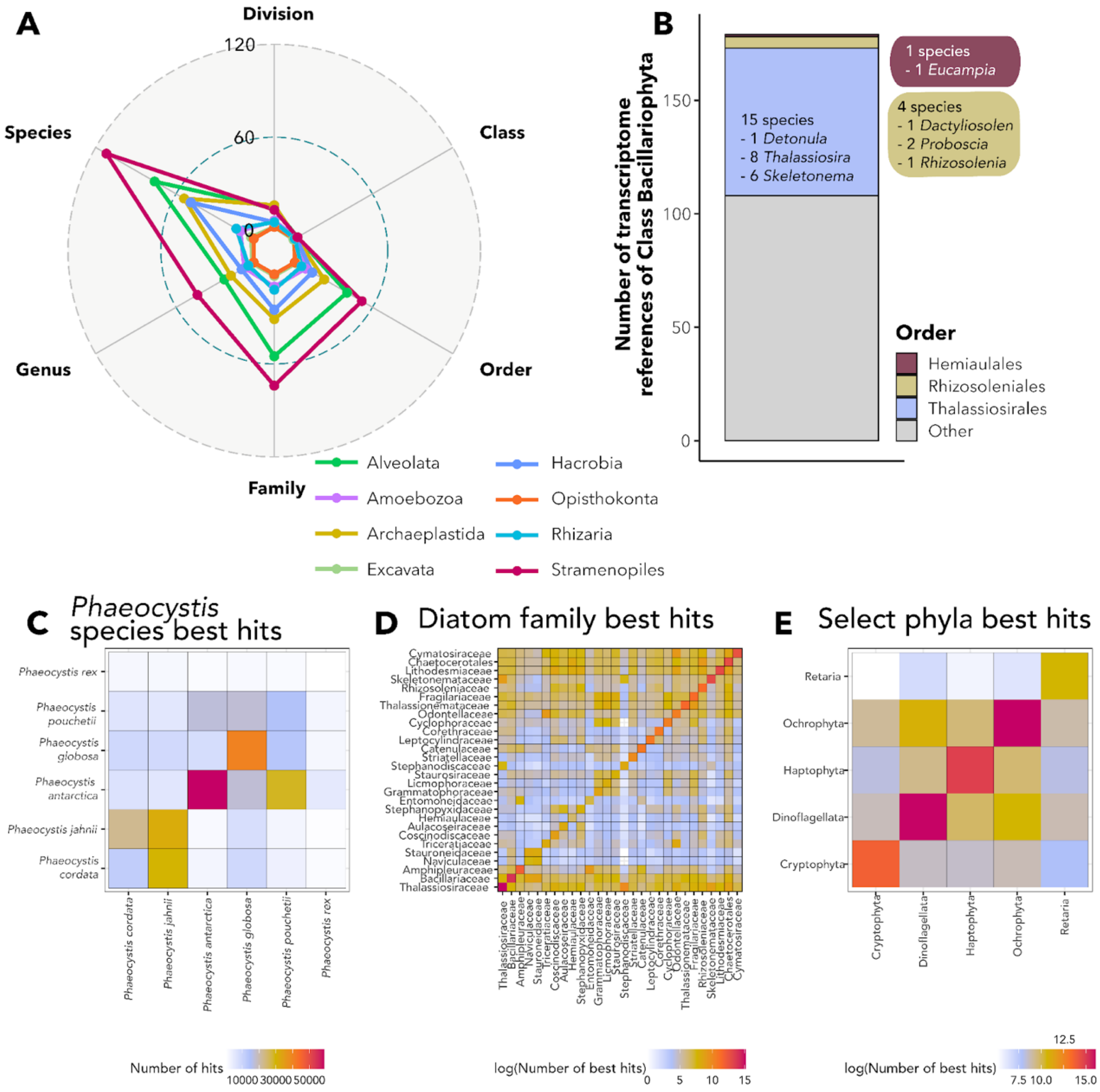


Supplementary Figure 2. Database imbalance in the Marine Microbial Eukaryote Transcriptome Sequencing Project (MMETSP) dataset. A: Number of available references at different levels of taxonomy (y-axes) for different major taxonomic groups of protists; B: Order-level breakdown of Families *Rhizosoleniaceae*, *Hemiaulaceae*, and *Skeletonemataceae*, three differentially-covered and differentially-annotated Families of diatoms in the Narragansett Bay study; C-E: Comparison of overlapping best BLAST hits at the three levels of taxonomic hierarchy in the paper. Some references contribute very few overlapping hits between sequences, indicating that these sequences will be less useful in recruiting best hits to sequences that are not near-exact replicates of the ones found in the database. For diatom families in D, many families have a similar or fewer number of top hits to other diatom families as compared to the family of which they are a member.

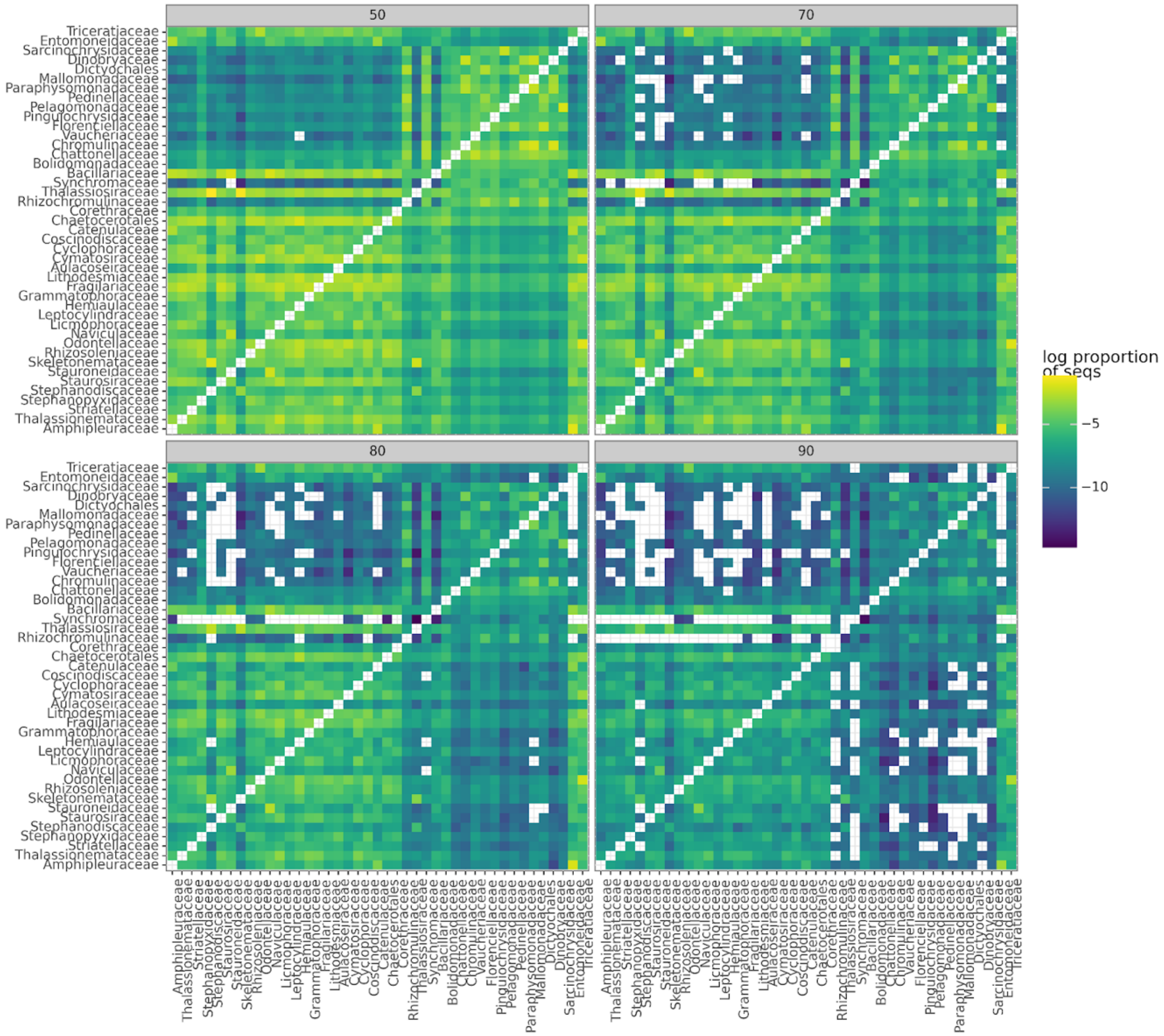


Supplementary Figure 3. Log-normalized proportion of sequences that have overlapping hits at a given (facets) percentage identity level among diatom families (within phylum Ochrophyta). Families are ordered according to hierarchical clusters of hits at the 90% sequence identity level.

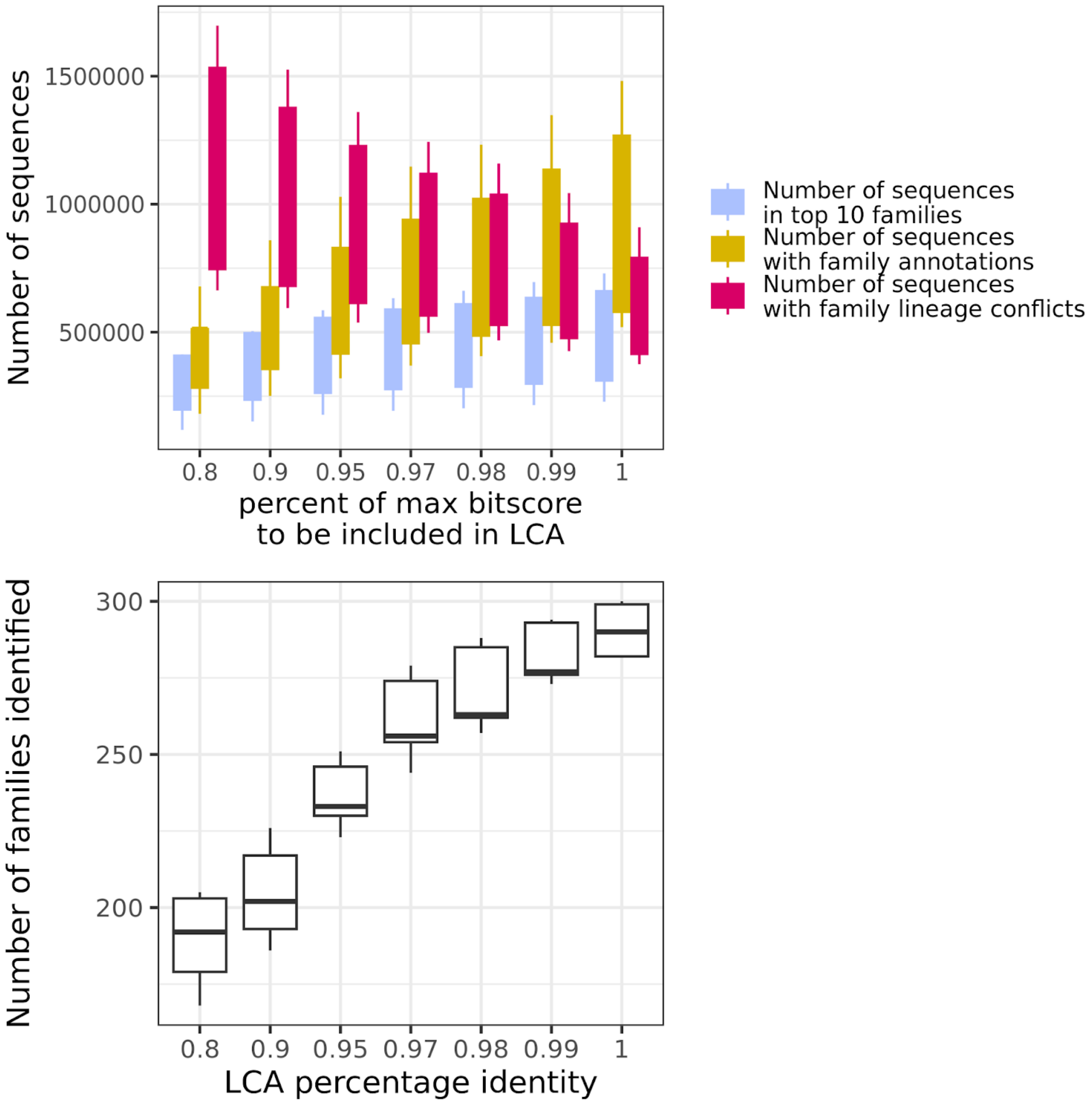


Supplementary Figure 4. Effect of setting the threshold for bitscores included in a least common ancestor search of Narragansett Bay sequences against the MMETSP + MarRef2 database. The value on the x-axis of each plot corresponds to the percentage of the maximum recorded bitscore that a hit would need in order to be considered in the search. Top: Effect of LCA bitscore threshold on the number of sequences annotated and sequences contained in the top 10 most abundant families. Bottom: Effect of the bitscore threshold on total families identified. The strong effect of modifying this parameter indicates that many hits with similar bitscores have different family annotations. Because not all hits have identical bitscore, adding new sequences to the database further complicates questions on how to interpret the quality of short alignments for taxonomic assignment.

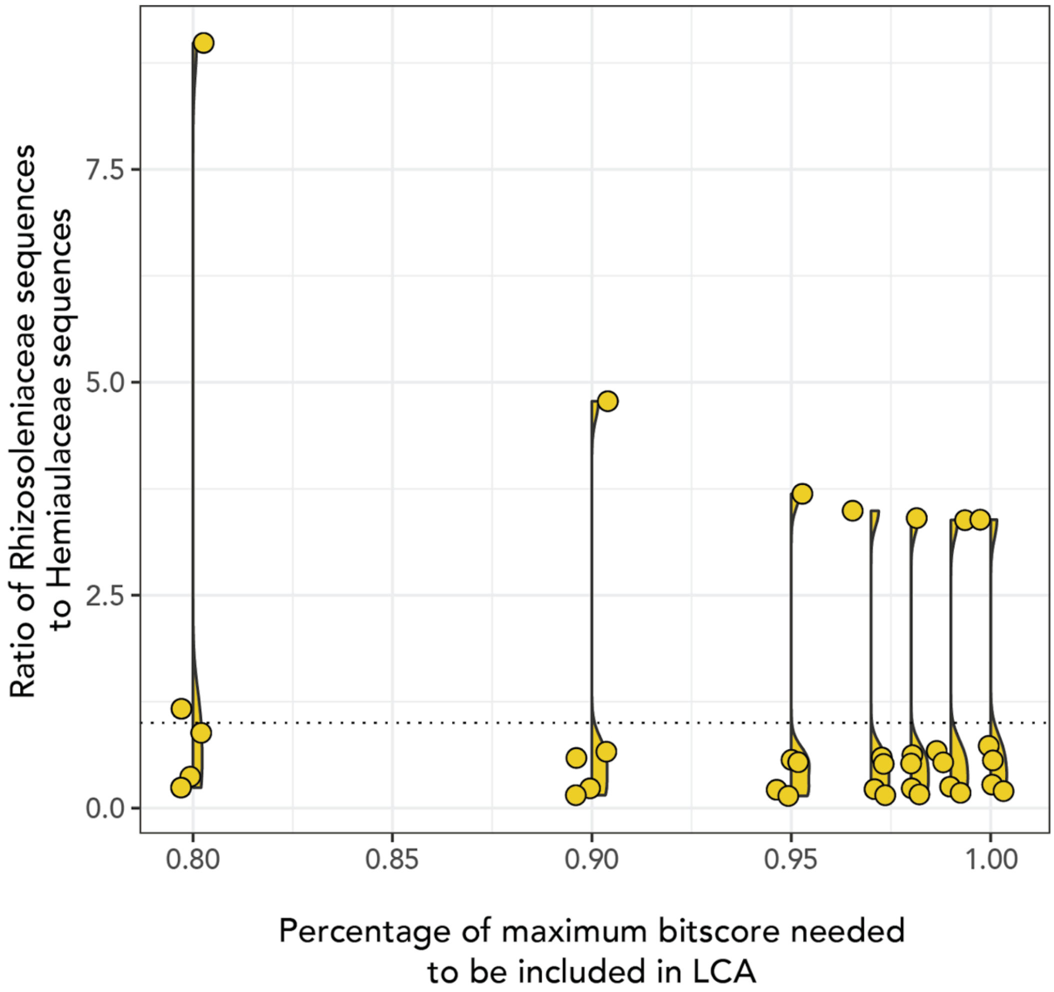


Supplementary Figure 5. Effect of the threshold used for a hit to be included in the LCA search on the ratio between family *Rhizosoleniaceae* and *Hemiaulaceae* sequences. The dotted horizontal line indicates the same composition of *Rhizosoleniaceae* and *Hemiaulaceae* sequences. Allowing more hits to be considered in the LCA can flip individual samples from *Hemiaulaceae* dominance to *Rhizosoleniaceae* dominance.

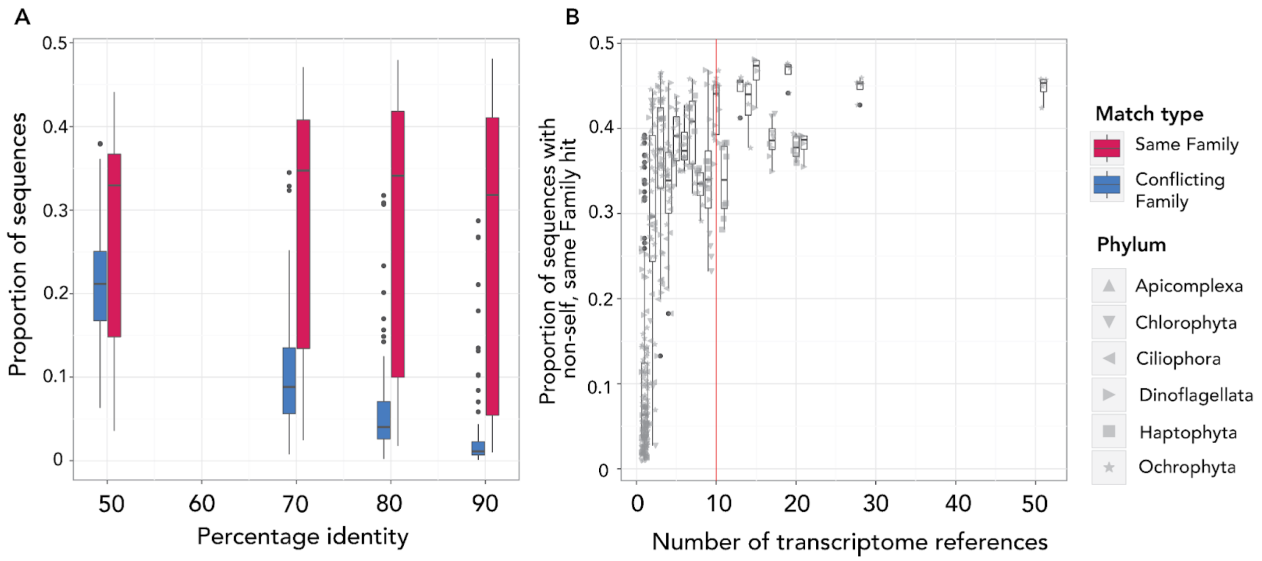


Supplementary Figure 6. Effect of database composition on effective annotation at the family-level. A: Proportion of MMETSP sequences with non-self hits to (right) the same family and (left) a different family using a bitscore cutoff of 50 and percentage identity cutoffs as indicated on the x-axis. While non-self hits to the same family does not change much on average with increasingly stringent percentage identity up to a cutoff value of 80%, the number of conflicts with inaccurate families drops with increasing percentage identity. B: Proportion of non-self target family hits made to families with the number of transcriptome references present in the database indicated on the x-axis. While families with very few references in the database tend not to have overlapping hits within the same family, the total number of sequences with same family hits tends to saturate just below 50% of total sequences regardless of the number of references added to the database. For some sequences, alignment-based annotation to the same family will be ineffective regardless of database size.

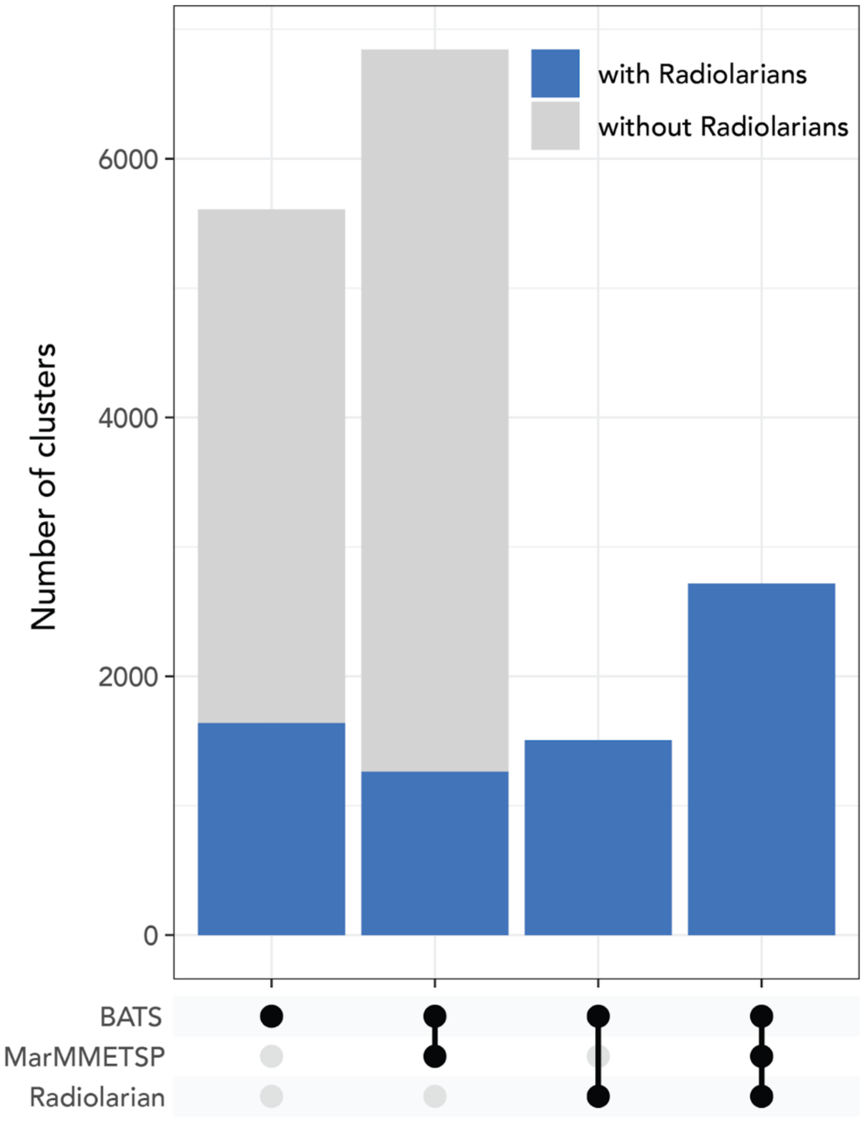


Supplementary Figure 7. Number of clusters formed by DIAMOND DeepClust on the BATS dataset before and after adding in radiolarian sequences to the original database (MarRef2 and MMETSP), shown for clusters that contain a sequence annotated to be radiolarian by the combined database. The overall number of clusters was reduced after adding in the radiolarian sequences, in addition to new sample sequences previously unclustered with any reference falling into clusters with radiolarian sequences.

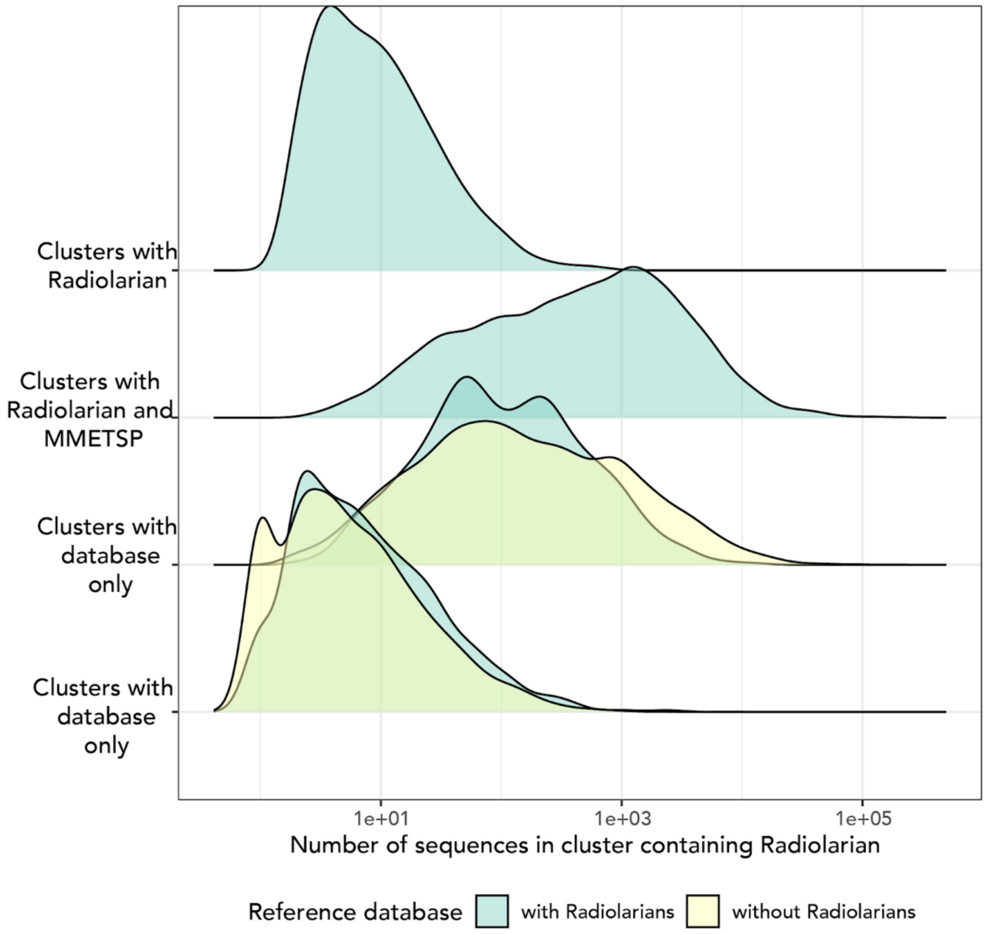


Supplementary Figure 8. Comparison of cluster sizes before and after adding Radiolarian sequences to a DIAMOND DeepClust search containing the MarRef2 + MMETSP database and the BATS dataset proteins, for clusters that contain sequences annotated to be Radiolarian by EUKulele. BATS protein clusters that contain both a Radiolarian sequence and a sequence from the MarRef2 + MMETSP database tended to be larger than the effect of adding the Radiolarians alone would suggest, indicating that expanding the database caused restructuring of clusters.

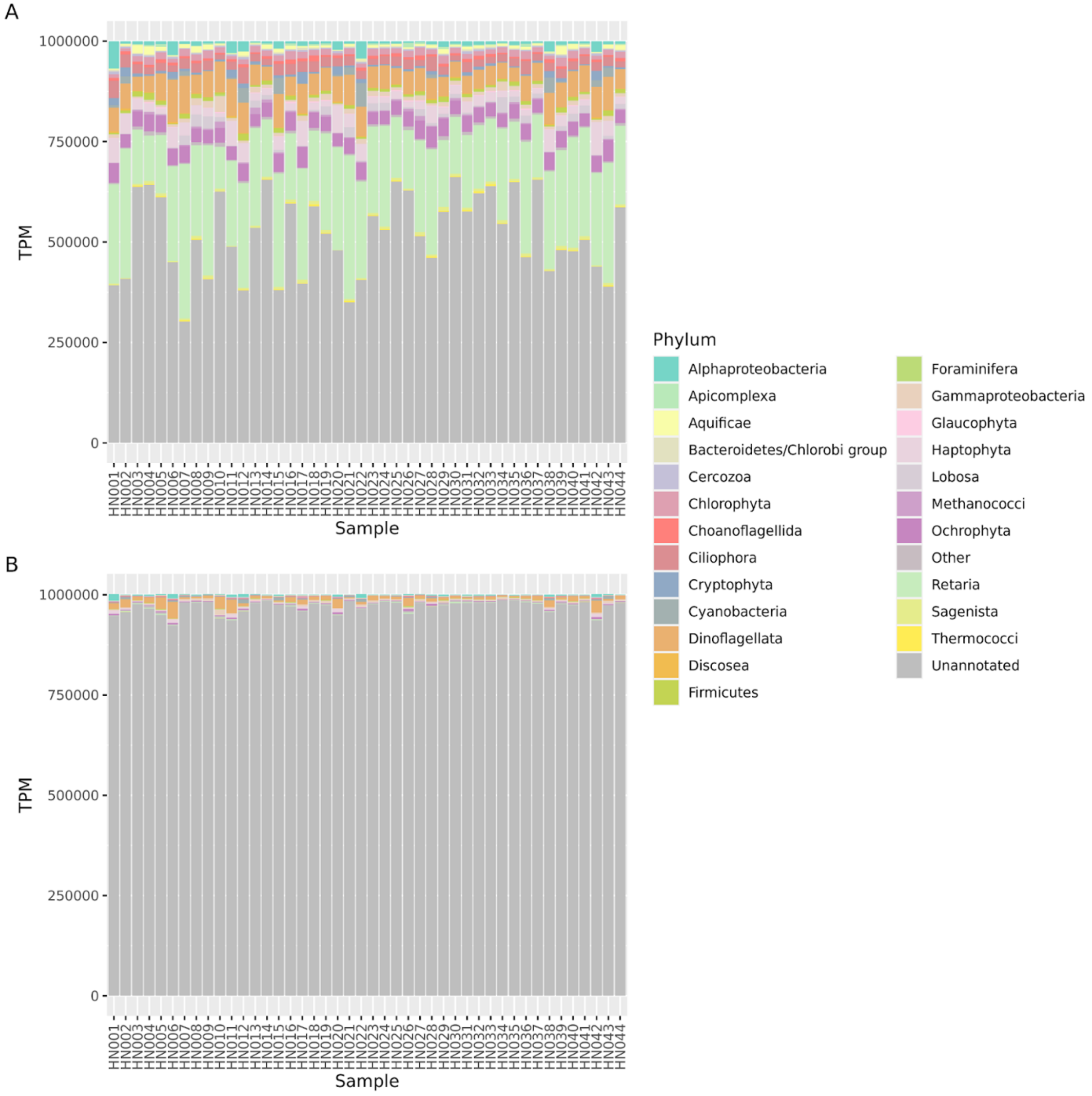


Supplementary Figure 9. BATS dataset showing comparison of percentage annotated across samples when the EUKulele mapping is done against ORFs (B) as compared to against contigs (A), even when the TPM is assigned in full to each predicted ORF (i.e. reads are mapped to ORFs). The dramatic difference between these two figures is partially due to the prevalence of diverse but offshore and often deeper below the surface samples in this dataset, with no closely related cultured representative available in databases. In the Narragansett Bay dataset, only 23.7% (1,806,887 of 7,624,296) predicted proteins were not assigned an annotation by EUKulele. In the BATS dataset, many fewer of the contigs which recruited reads contained a high quality (>300 bp) predicted protein in the first place, but many of these contigs contained regions which could be aligned to the protein database by EUKulele (A).

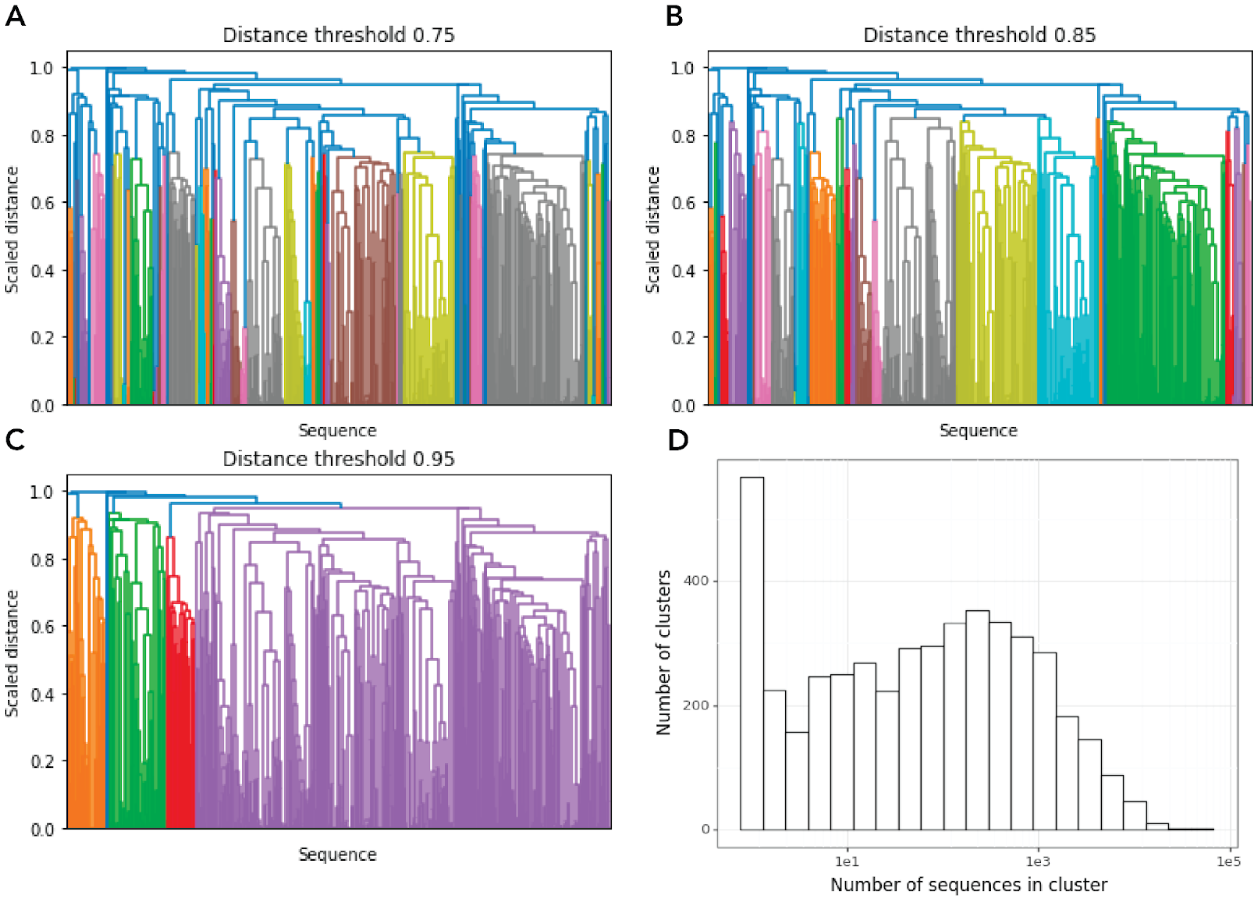


Supplementary Figure 10. A-C: effect of clustering threshold choice on number of clusters generated by hierarchical clustering for one randomly chosen homology cluster. D: for the 5,000 sequences randomly selected from the MMETSP and MarRef, the number of sequences contained in DIAMOND DeepClust [23] clusters containing that sequence. While 567 sequences were contained in clusters only containing one sequence, most co-clustered with other database references. By comparison, 1,388 of these sequences were not annotated at the phylum-level by EUKulele, and 82 were misclassified at this level.

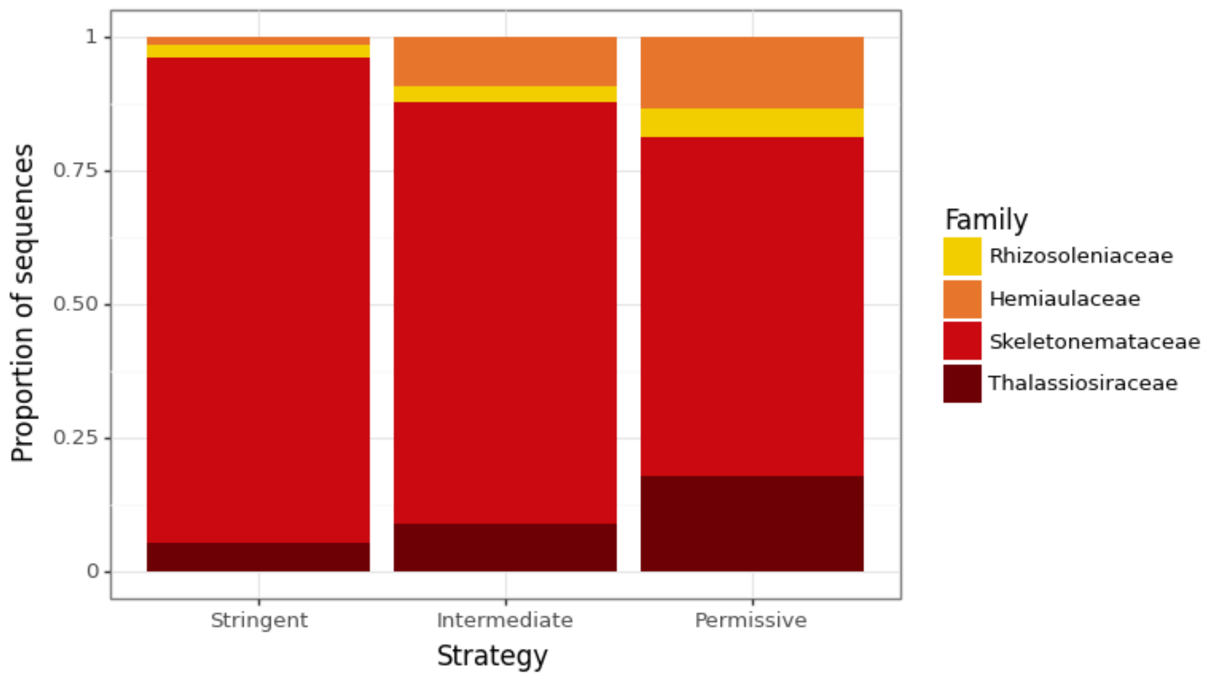


Supplementary Figure 11. Relative abundance of sequences from each of the four considered diatom families using the three clustering strategies.

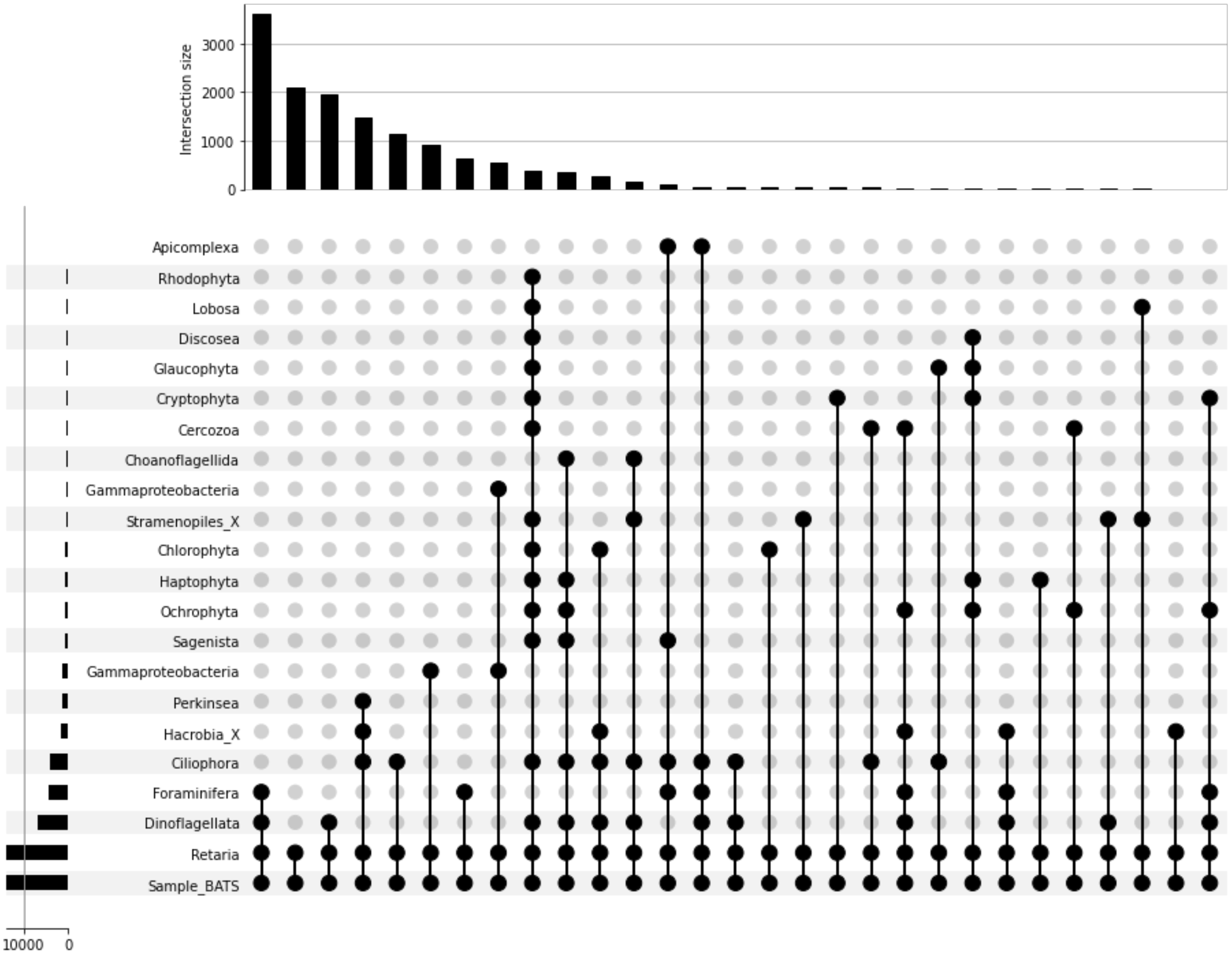


Supplementary Figure 12. Division breakdown of clusters containing radiolarians (Retaria in this figure and in our taxonomy table; note that strictly Foraminifera and Radiolaria are both infra/subphya of Retaria) using the stringent strategy. The UpSet plot shows the number of BATS sequences found in clusters with major eukaryotic groups. The bars arranged horizontally across the top of the plot show the number of BATS sequences that fall into each category, arranged from most to least common. The bars arranged vertically indicate the size of the sequence dataset represented in the plot. The majority of sequences from the BATS dataset fall into clusters that contain radiolarians also contain sequences from foraminifera and dinoflagellates, even using the stringent strategy.

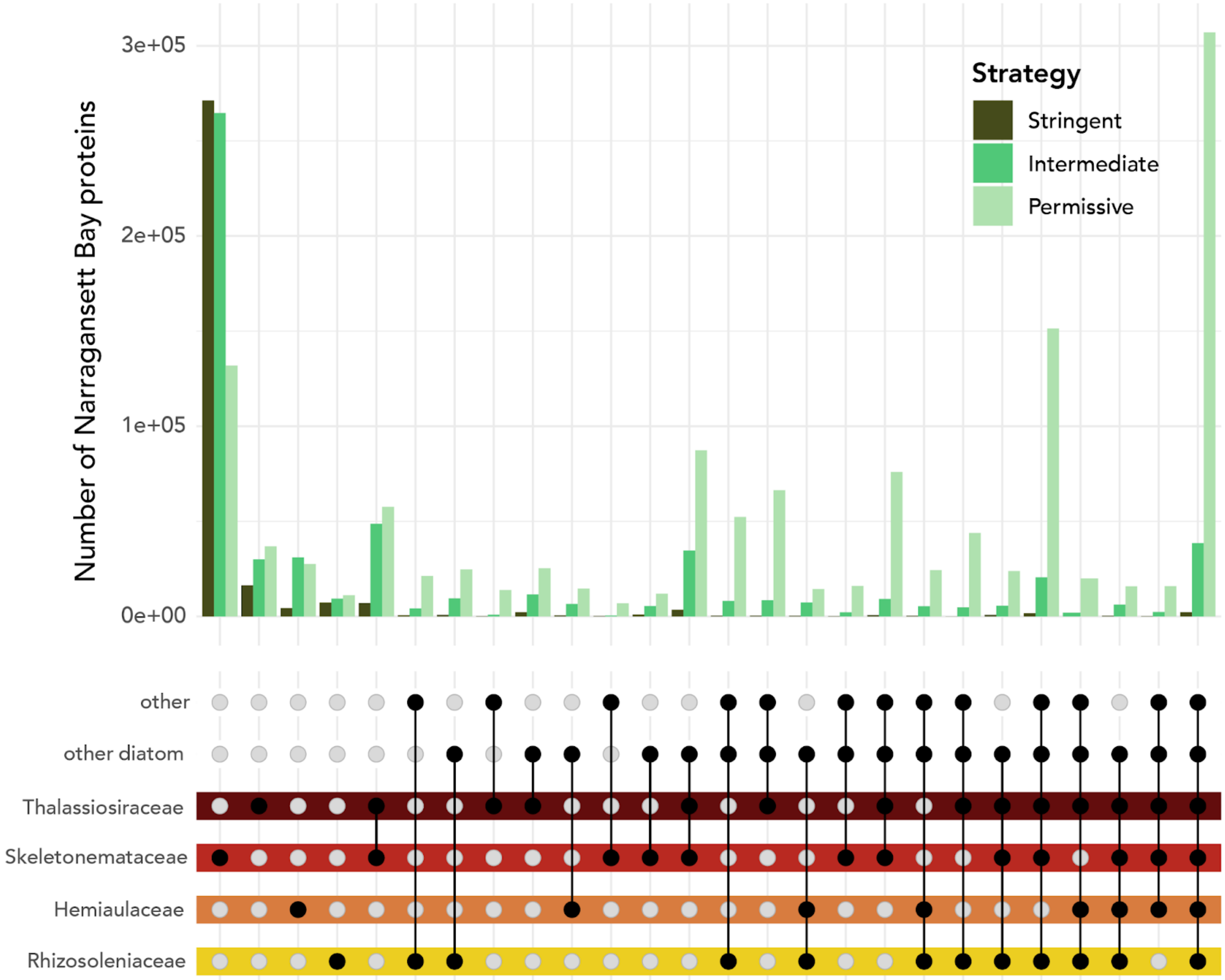


Supplementary Figure 13. Tax-aliquots clustering demonstrates relatedness between diatom families and annotation confidence. Upset plot showing the number of Narragansett Bay sequences found in clusters with some combination of diatom taxa. The color of each bar corresponds to the strategy used by the clustering run, from permissive (clusters are based on less putative relatedness) to stringent (high relatedness is needed for a cluster to be retained).

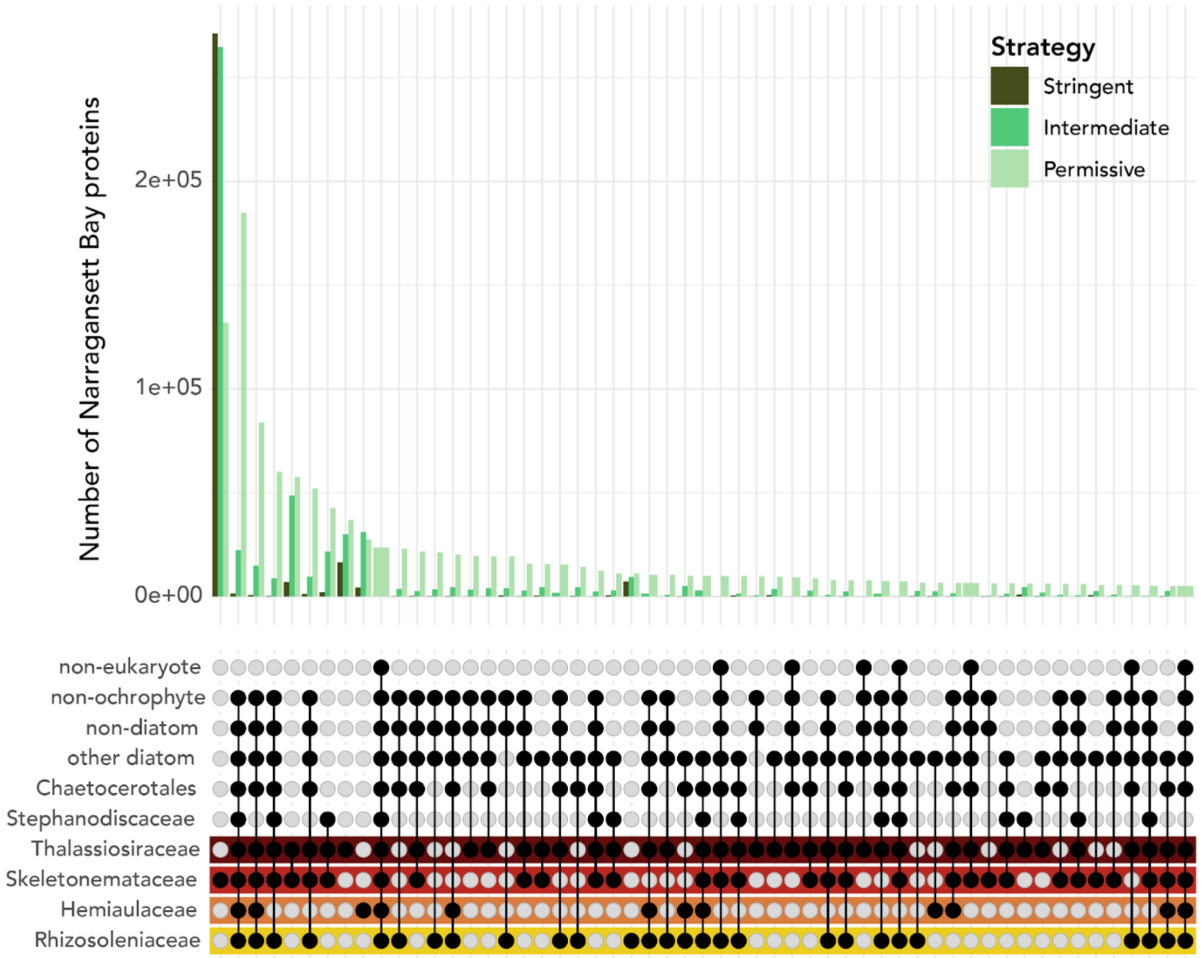


Supplementary Figure 14. Identical to Supplementary Figure 13, but with higher resolution of non-target groups. Tax-aliquots clustering demonstrates relatedness between diatom families and annotation confidence. Upset plot showing the number of Narragansett Bay sequences found in clusters with some combination of diatom taxa. The color of each bar corresponds to the strategy used by the clustering run, from permissive (clusters are based on less putative relatedness) to stringent (high relatedness is needed for a cluster to be retained).
